# Alkamines reveal a hidden layer of steroid and drug metabolism

**DOI:** 10.64898/2026.05.13.724743

**Authors:** Julius Agongo, Subhaskar R Panga, Shipei Xing, Vincent Charron-Lamoureux, Harsha Gouda, Yasin El Abiead, Megan R. Nelson, Abubaker Patan, Marvic Carrillo Terrazas, Kine Eide Kvitne, Jeong In Seo, Prajit Rajkumar, Sadie Giddings, Helena Mannochio-Russo, Jasmine Zemlin, Ipsita Mohanty, Marta Sala-Climent, Zhewen Hu, Victoria Delaray, Simpa K. Yeboah, Haoqi Nina Zhao, Andrés M. Caraballo-Rodríguez, Claire E. Williams, Candace L. Williams, Wilhan D. Gonçalves Nunes, Kathleen Dorrestein, Jennifer Cao, Isabelle Shepherd, Rachel Bock, Nathaniel Roethler, Adrian Jinich, Lindsey A. Burnett, Jeremy Carver, Robert N. Devine, Christopher K. Arnatt, Iain A Murray, Rob Knight, Monica Guma, Lee R Hagey, Gary Perdew, Nuno Bandeira, Mingxun Wang, Hiutung Chu, Dionicio Siegel, Pieter C. Dorrestein

## Abstract

Biomedical research overlooks most genes in favor of a well-studied minority, yet whether analogous blind spots exist in metabolomics remains unknown. We show that reductive amination, forming secondary amines from aldehydes or ketones and amines, generates a previously hidden class of metabolites we term alkamines. Multiplexed synthesis of 8,475 alkamines combined with MS/MS searches across 1.7 billion spectra identified 1,626 candidates across multiple species and organs. Of these, 56 were confirmed in biological samples, including 27 steroid- and 12 drug-derived alkamines matching prescription patterns. Notably, 77% of synthesized alkamines are absent from PubChem. This combinatorial logic likely explains why alkamines have evaded detection and suggests drug metabolism frameworks substantially underestimate drug-derived metabolite diversity. Reductive amination is an overlooked route modifying steroids, bile acids, and xenobiotics.

## Introduction

In genomics, fewer than 10% of genes account for approximately 90% of publications(*1*). This imbalance reflects historical patterns of experimental preference rather than biological or clinical importance. Importantly, understudied genes are not missing from experimental datasets. Rather, they are present, but are often excluded before results are reported. The bias arises not from the biology itself, but from choices in sampling, detection, annotation, and reporting. We asked whether an analogous blind spot exists in metabolomics. Even in 2026, could entire classes of detectable molecules remain undiscovered, not because they are rare or chemically inaccessible, but because they have simply not been examined in existing biological data?

Metabolism shapes the chemical landscape of life, yet the full repertoire of biologically manifest transformations remains incompletely mapped. In a preprint it has been reported that in untargeted metabolomics datasets from one of the most extensively studied sample matrices, human serum, a mass spectrometry feature detected in more than 50% of samples has only about a 50% chance of being annotated (*2*). When detection frequency drops to 10% of samples, the probability of annotation falls to ∼5%, and declines even further for rarer features. For less-studied sample types, annotation rates are substantially lower. The biochemical transformations most likely to evade current annotation approaches are those generated combinatorially through interactions among dietary components, host physiology, microbiome composition, enzymatic activities, and cofactor availability. The combinatorial nature of these processes may generate not just a single molecule but extensive chemical diversity, producing molecular families that are collectively abundant even when individual compounds occur only sporadically across samples. We therefore hypothesized that hidden classes of molecules generated through such combinatorial reactions are already present in public metabolomics data but remain unannotated.

Among the most fundamental reactions in organic chemistry, reductive amination converts aldehydes or ketones and amines into substituted secondary amines through imine/imminium intermediates (**Fig. 1A**). Genomic and metagenomic mining has established that imine reductases are broadly distributed among living organisms, including but not limited to bacteria, fungi, plants, protists and animals, with systematic surveys identifying hundreds of sequences that are consistent with enzymes that would have imine reductase activities(*3, 4*). Over 300 sequences of diverse enzymes have been shown to be capable of catalyzing this chemistry(*5, 6*). The panel of enzymes uncovered with this activity is distributed across the diversity of cellular life, from bacteria to eukaryotes, including human imine reductases. Despite the ubiquity of all the required functional groups and enzymes in biology, the hypothetical products of these reactions have remained almost entirely absent from our understanding of human and animal metabolism(*7, 8*). We additionally hypothesized that this absence is not due to a lack of chemical reactivity, but instead due to an annotation gap.

**Fig. 1.**
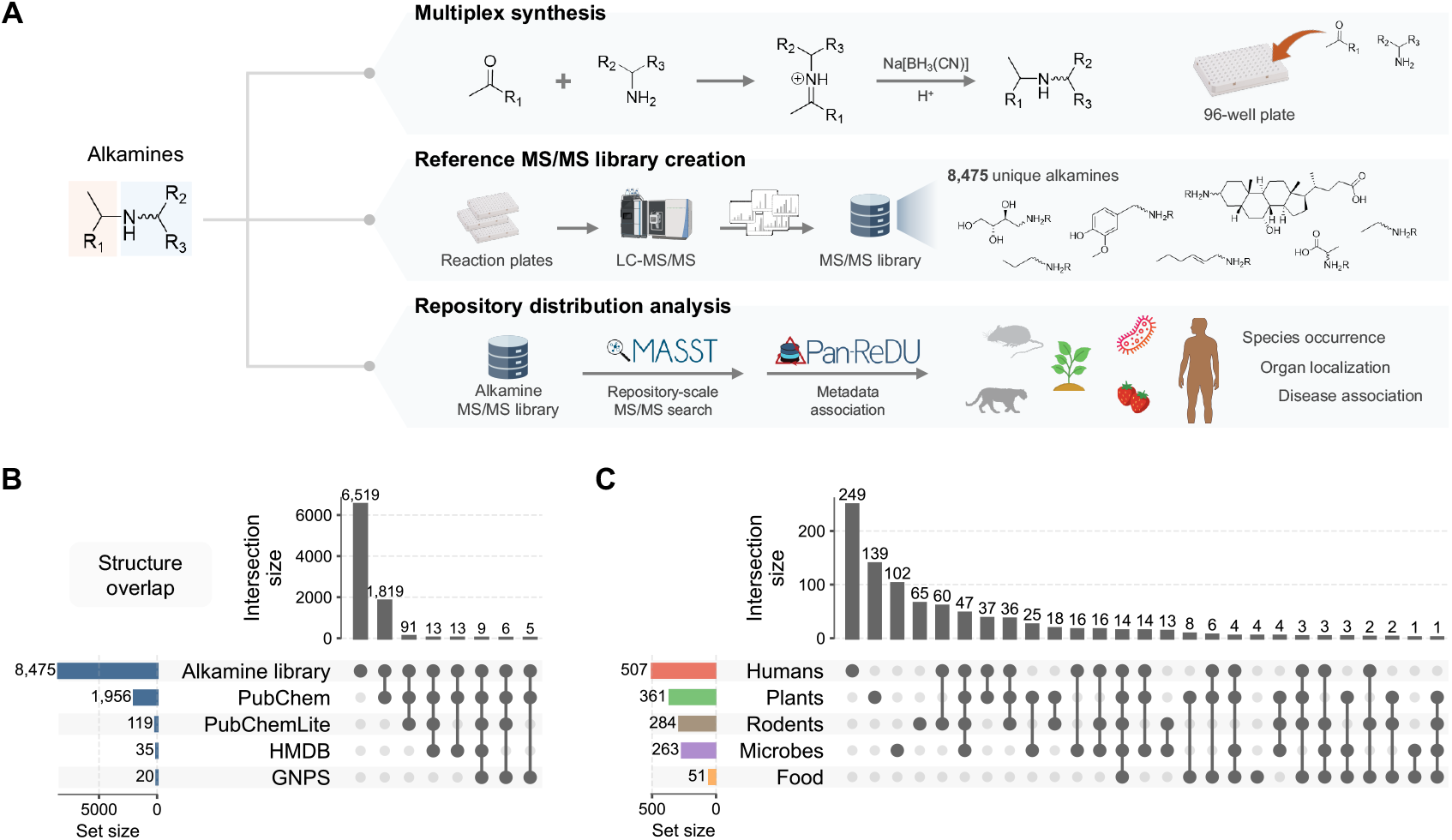
Overview of alkamines created using synthetic multiplexing and reverse metabolomics. **(A)** Schematic of alkamine definition and multiplexed synthetic workflow. Structurally diverse aldehydes and ketones were reacted with different amines to generate 8,475 unique alkamines using multiplexed synthesis, which are then subjected to reverse metabolomics analysis. (**B)** Summary of chemical diversity captured in the synthetic library. **C**, Summary of sample type metadata distribution of MS/MS matches to alkamines to public LC-MS/MS metabolomics datasets, highlighting widespread occurrence across humans, rodents, plants, and microbes of the MS/MS matches belonging to this metabolite class. All icons were obtained from bioart, bioicons website.

The scale of what theoretically and combinatorially exists is striking. KEGG and similar metabolic pathway maps describe biological systems as containing thousands of aldehydes, ketones, and amines that in principle could construct millions of structurally distinct alkamines(*9*). Indeed, a recent preprint reported the reductive amination product for methylglyoxal and taurine(*10*). In plants and marine invertebrates such as mollusks, secondary amines formed through reductive amination, known as opines, are characterized(*11, 12*). Sharks produce the aminosterol squalamine, a secondary amine also presumably generated through reductive amination(*13*). Yet in humans and other mammals, saccharopine and pipecolate, both intermediates of lysine metabolism, and glutamate-5-semialdehyde, a precursor to proline, represent the only secondary amines arising from reductive amination that we could find documented in humans(*14*–*16*). Structurally *N*-alkylated amino acids and related products, despite the well-documented abundance of suitable carbonyl precursors reaching millimolar concentrations in human biofluids, have not been reported.

The oversight has a clear cause. In LC-MS/MS-based discovery metabolomics, molecule identification depends on matching MS/MS spectra to reference libraries(*17, 18*). No such library existed for reductive amination products, molecules that are individually rare, yet arise from enzymatic machinery common across biology, and whose formation depends on the simultaneous availability of specific substrates, co-factors, and enzymes. Without reference spectra, annotations cannot be made, and entire classes of metabolites remain outside the annotatable metabolome, even if they are abundant. The recent growth of large public metabolomics repositories, now containing over 1.7 billion spectra from thousands of datasets, combined with fast repository-scale search engines, creates for the first time the analytical infrastructure needed to systematically address this gap(*19, 20*).

In sequencing, strategies for identifying missing rare gene annotations have been developed by reconstructing plausible ancestral state sequences(*21, 22*); however, this concept has not been extended to metabolomics, where entire classes of metabolites likely exist that have been detected, yet remain unannotated. Here, we describe a synthesis-driven reverse metabolomics strategy to allow identification of reductive amination products. Aldehydes and ketones (96 of both) were combined with 110 amines using multiplexed synthesis to generate an MS/MS library of 8,475 alkamines. This library was used in search across public repositories spanning humans, animals, plants, and microbes(*23, 24*). This pan-repository screening approach identified 1,626 candidate alkamines, of which 56 were confirmed in biological samples by retention time and 20 by additional ion mobility matching. Notably, we find that two intensively studied molecular classes in biology, steroids and drugs, undergo reductive amination to generate previously unknown metabolites that are commonly observed in mass spectrometry data of biological samples. These findings reveal a gap in how steroid and xenobiotic metabolism has been characterized.

## Results

### MS/MS of alkamines are widely observed in public LC-MS/MS data

To obtain MS/MS spectra of alkamines, 96 structurally diverse aldehydes and ketones were reacted with 110 amine-containing compounds via reductive amination using cyanoborohydride, the approximate organic chemical equivalent of NAD(P)H in enzyme catalyzed reduction reactions(*25*) (**Fig. 1A**). Unlike enzyme-based reductions, when one of the amines or the carbonyl containing compound is chiral, this reaction generates diastereomeric secondary amines with low to no selectivity(*26*). Amines were grouped into batches of 20 and reacted with each carbonyl in a 96-well plate format, enabling multiplexed synthesis. Alkamine products were subsequently linked to their corresponding MS/MS spectra by matching LC-MS/MS scan numbers from the multiplexed reactions to predicted product SMILES(*27*) (see Methods for details). In total, 18,031 MS/MS spectra were linked to 8,475 unique alkamines, including 15,023 [M+H]^+^, 2,857 [M+Na]^+^, and 151 [M+NH_4_]^+^ ion species (**Table S1**). Notably, 77% of the 8,475 synthesized alkamines are not documented in PubChem, one of the most comprehensive structural databases in existence (**Fig. 1B, Table S2**).

Reverse metabolomics searches across four public repositories (MetaboLights, Metabolomics Workbench, NORMAN digital sample freezing platform, and GNPS/MassIVE, indexed September 2025) yielded 418,368 MS/MS matches at a cosine similarity of 0.8 or greater, a threshold associated with false discovery rates below 1% for MS/MS matching(*19, 28*). These matches represent level 2/3 annotations according to both the Schymanski levels and Metabolomics Standards Initiative frameworks, constituting the highest confidence achievable without access to the original biological specimens(*29, 30*). As with any MS/MS library-based annotation, matches may represent different ion forms, structural isomers, or products of in-droplet electrospray reactions rather than pre-existing biological metabolites(*31*). Level 1 identification requires additional matching of chromatographic retention time and, where available, ion mobility drift time against authentic synthetic standards, an approach we apply to a subset of compounds (see examples below). Across the library, 1,626 of 8,475 synthesized alkamines had at least one repository MS/MS match (**Table S3**).

Using harmonized sample metadata from PanReDU for human and rodent datasets, and domain-specific MASST searches for microbial monocultures, plants, and foods, we evaluated the organismal, organ, and condition-specific distributions of matched alkamines. MS/MS matches were obtained for 507 alkamines in human, 361 in plant, 284 in rodent, and 263 in microbial LC-MS/MS data, with 60 alkamine annotations overlapping between human and rodent datasets(*23*) (**Fig. 1C, Table S4**). The most frequently matched human alkamines were synthesized from phenylalanine with 2-hydroxypropanal and GABA with pyruvaldehyde, with 4,173 and 5,464 MS/MS matches, respectively. Alkamines overlapping between human and microbial datasets were predominantly derived from amino acids and amino acid-derived amines, including tryptamine and GABA, paired with short-chain carbonyls, including pyruvaldehyde, propenaldehyde, butyraldehyde, and indole-3-pyruvic acid, suggesting a potential host-microbe axis in alkamine biosynthesis. Fourteen alkamines had MS/MS matches across all five biological kingdoms (**Fig. 1C**).

Considering all MS/MS matches across the full alkamine library, reverse metabolomics provided 1,626 matched compounds arising from reductive amination across 110 amines and 96 aldehydes or ketones (**Fig. 2**). Matched alkamines were derived from diverse amine classes, including proteinogenic and non-proteinogenic amino acids, modified amino acids, polyamines, neurotransmitters, peptides, biogenic amines, drugs, and nucleic acid-derived compounds. These annotations are level 2, a structural hypothesis, and should be interpreted accordingly. Level 1 identification requires matching both MS/MS spectra and chromatographic retention time from biological extracts against standards. This approach provides additional support for its molecular identity. We apply this approach to a subset of alkamines below.

**Fig. 2.**
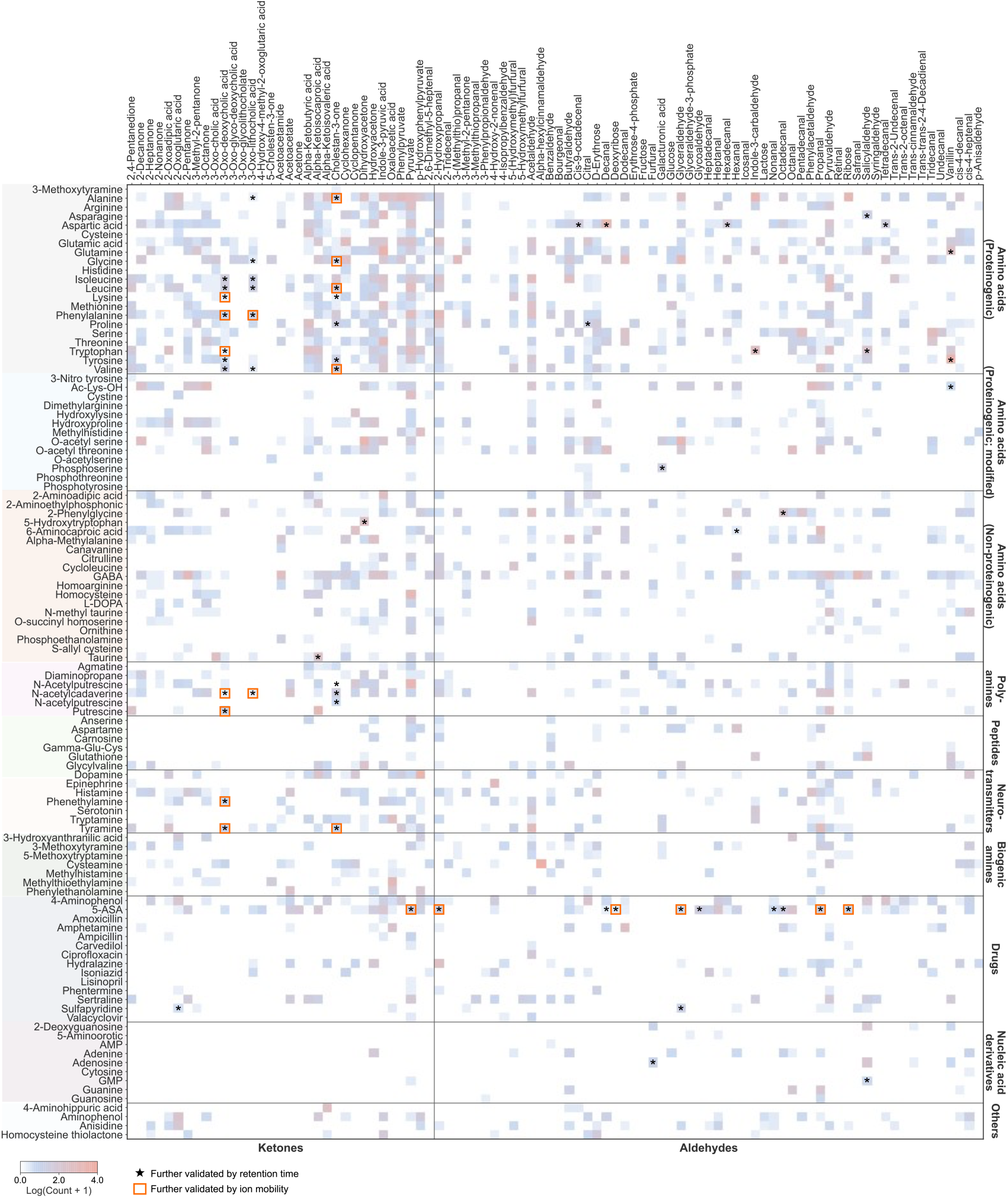
Heatmap of MS/MS matches from 1,626 alkamines across public datasets. The heatmap shows MS/MS matches of alkamines to pan-repository public MS/MS data using the synthesized multiplexed alkamine library. Black stars indicate matches supported by retention time and MS/MS comparison between synthetic standards and the biological sample. Red squares indicate matches further supported by ion mobility-based collision cross-section measurements. Validation by chromatography coelution of the MS/MS matches was done based on the availability of the biological samples.

### Reductive amination reveals a hidden layer of steroid metabolism

Bile acids and other cholesterol-derived steroids are among the most extensively studied metabolite families, yet their chemical diversity has continued to expand with the discovery of previously unrecognized microbial and host transformations(*32, 33*). We asked whether ketone-containing bile acids and related steroids could undergo reductive amination to form alkamines (**Fig. 4A**).

To obtain additional evidence supporting structural hypotheses for a subset of candidate alkamines, we identified available samples or comparable matrices from the MS/MS matches to public data likely to contain target compounds based on FASST search results. We selected representative examples spanning each amine class, including drug-derived, neurotransmitter-derived, and amino acid-derived alkamines, according to sample availability. We then performed LC-MS/MS analysis of these samples alongside authentic synthetic standards, matching MS/MS spectra and chromatographic retention times. In total, 56 compounds achieved Level 1 identification by MS/MS and retention time matching in human, lizard and feline fecal samples, including bile acid, cholestane, and 5-ASA alkamine conjugates, 20 of which were further supported by ion mobility drift time matching (**Fig. S4-S7; Table S5**).

Examining the distribution of alkamine MS/MS matches across sample metadata in PanReDU, feces, urine, and blood exhibited the highest diversity of matched alkamines in both humans and rodents (**Fig. 3A,B**), consistent with the metabolomic coverage of these sample types in public repositories. Notably, the human brain and breastmilk also contained substantial numbers of MS/MS matches despite being comparatively rare in public datasets, suggesting alkamine formation extends to tissues beyond the gut and circulation. Across all sample types, alkamines derived from tyrosine, dopamine, alanine, cysteamine, hydroxyphenylpyruvate, indole-pyruvic acid, pyruvate, and vanillin were detected across the largest number of data files and biological matrices.

**Fig. 3.**
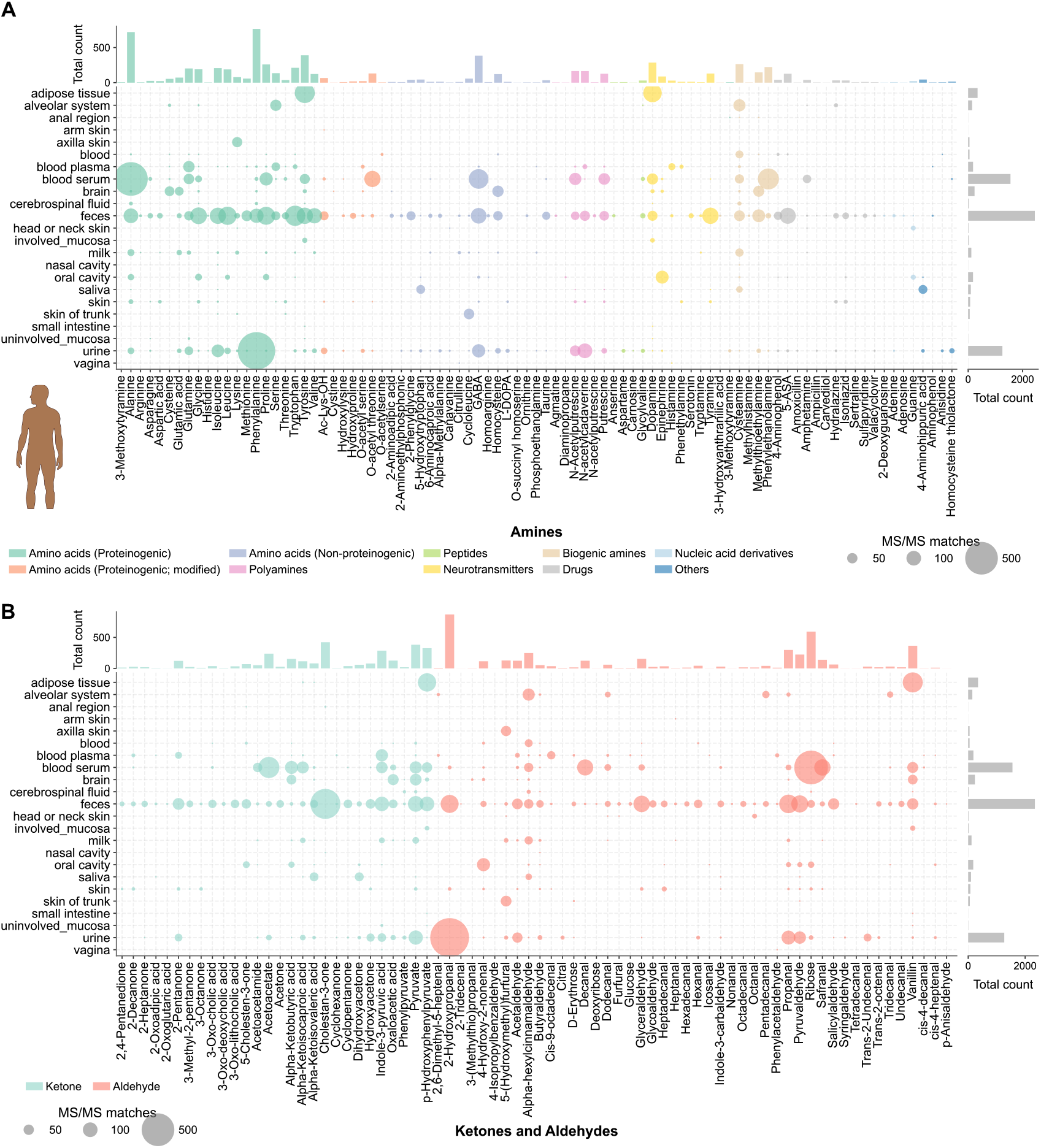
Pan-repository MS/MS matches and their organ and biofluid distributions in humans. **(A)** Distribution of MS/MS of alkamines synthesized from amines in human samples. **(B)** Organ and biofluid distribution of MS/MS of alkamines synthesized from aldehydes and ketones in human samples. The same data but normalized to sample number can be found in the supporting information (**Fig. S3A**,**B**).

As part of the multiplexed alkamine library, we synthesized alkamine derivatives from five ketone-containing bile acids and 35 amines. Because bile acids also undergo amidation at the terminal carboxylate(*34*), which proceeds through distinct chemistry from alkamine formation but could produce structurally similar products, there was potential for MS/MS spectral overlap leading to incorrect annotations (**Fig. 4B**). Several observations supported alkamines at the steroid scaffold rather than amidates for the MS/MS matches with the alkamine library. First, MS/MS spectra were detected for bile acids in which the carboxylate was already blocked by conjugation, as well as matched to steroids lacking a carboxylate entirely that cannot form amidates. Direct comparison of reference spectra showed that alkamine and amidate products produce readily distinguishable fragmentation patterns. Alkamine formation at the C3 position yielded much intense diagnostic fragment ions at *m/z* 359.294, 357.278, and 355.260 for lithocholic acid, deoxycholic acid, and cholic acid, respectively, consistent with secondary amine formation at the C3 position of the steroid core (**Fig. 4B**). A MassQL(*35*) query based on these diagnostic ions was applied to ensure that all reported MS/MS matches exhibited this characteristic fragmentation pattern. No amidate matches were found. Beyond bile acids, the multiplexed alkamine libraries contained five additional cholesterol-derived steroidal ketones: 5-cholestan-3-one, cholestan-3-one, estrone, tetrahydrocortisol, and pregnanolone (**Fig. 4A**). Pan-repository searches revealed MS/MS matches to bile acid and steroidal alkamines across multiple carbonyl-containing scaffolds and amine classes, with MS/MS of 91 amine-ketosteroid or keto bile acid pairs spectral matches in total, indicating that reductive amination occurs across diverse steroid backbones rather than reflecting a single substrate-amine pairing (**Fig. 4C**).

**Fig. 4.**
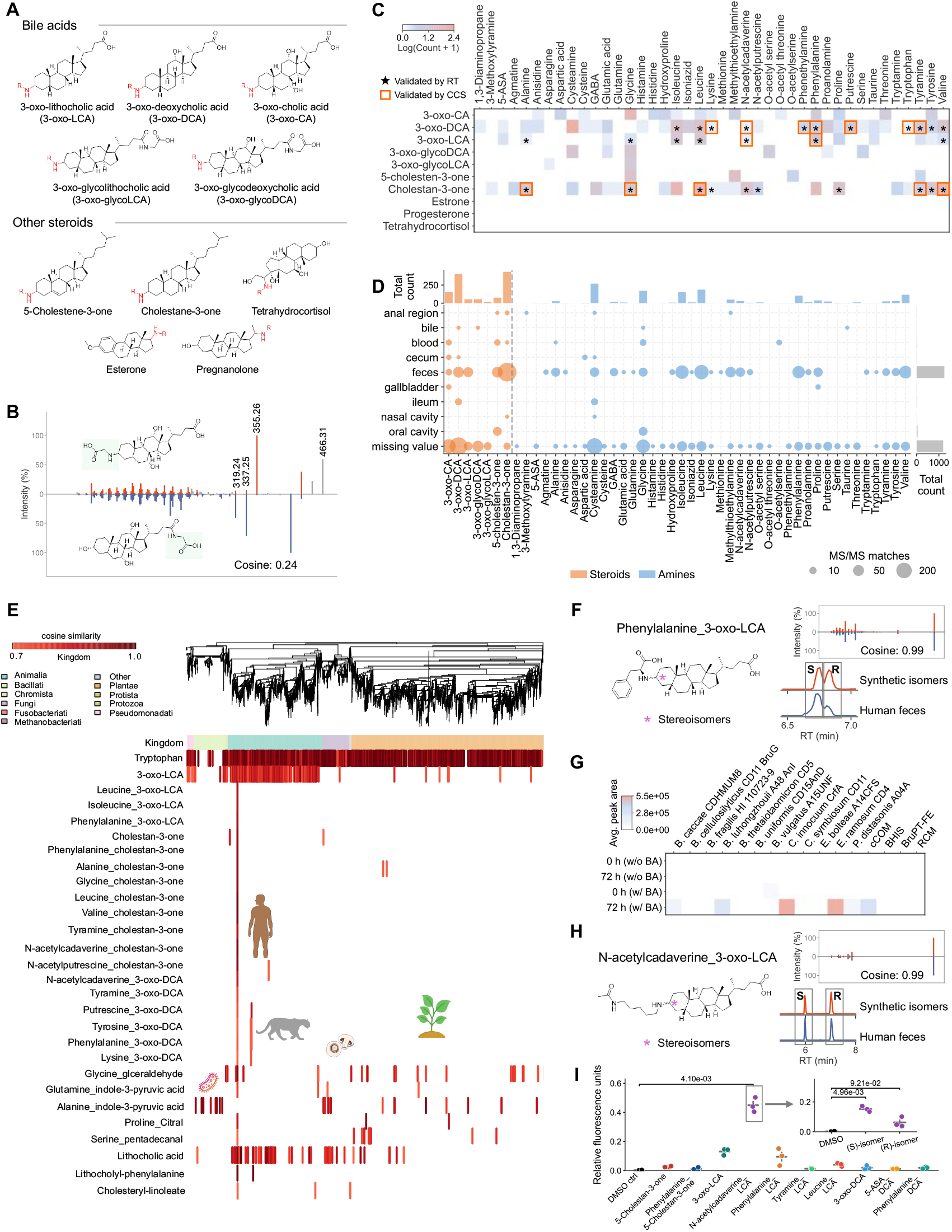
Reductive amination diversifies cholesterol-derived steroids, including bile acids. **(A)** Core structures of ketone-containing bile acids and cholesterol-derived steroids used for multiplexed alkamine synthesis. **(B)** Mirror plot comparing representative MS/MS spectra of alkamine and amidate derivatives, illustrating distinct fragmentation patterns that differentiate ketone modification from carboxylate amidation at the same collision energy. The cosine between these two spectra is very low at 0.24. **(C)** Alkamine MS/MS matches across bile acid (and steroidal carbonyl substrates (y-axis) and amines (x-axis) detected in public metabolomics datasets. **(D)** Organ, tissue and biofluid distribution of MS/MS derived from steroidal alkamines across public datasets. **(E)** Taxonomic distribution of MS/MS of bile acid and steroidal alkamines across public metabolomics datasets. MS/MS matches to tryptophan are shown as representative of a very commonly detectable metabolite that should be detectable in most organisms. **(F)** MS/MS and retention time matching of synthesized Phenylalanine_3-oxo-LCA alkamine to human feces. **(G)** microbial-mediated formation of Phenylalanine_3-oxo-LCA in 12-member single strain microbes and combined community (cCOM) in the presence of Reinforced Clostridial media (RCM), Brucella with pyruvate, taurine, and iron (BruPT-FE), brain heart infusion supplemented (BHIS) under anaerobic conditions (10% CO2, 7.5% H2, and 82.5% N2) (BHIS). **(H)** HL-60 calcium mobilization cell-based assay for broad screen of steroid alkamines towards GPCR and nuclear receptor binding, *N*-acetyl-cadaverine-LCA alkamine (isomeric mixture). Insert **(I)** is testing the S and R isomer of *N*-acetyl-cadaverine-LCA alkamine after purification.

MS/MS of bile acid and steroidal alkamines were most commonly detected in fecal samples but were also observed in intestinal tissues, bile, gallbladder, and blood, indicating both local and systemic occurrence (**Fig. 4D**). Taxonomic analysis of matches to public data using PanReDU harmonized metadata revealed that steroidal alkamines displayed strong animal-specific distributions(*36, 37*). MS/MS of cholestan-3-one derived alkamines were detected in 314 of 660,127 animal-associated files versus 5 of 226,312 non-animal files (Odds ratio [OR] = 21.54; 95% CI 8.9 to 52.1; Fisher’s exact test, p = 4.22 × 10^−33^), while MS/MS of 3-oxo-lithocholic acid and 3-oxo-deoxycholic acid alkamines were associated with animal files, with 34 of 660,127 versus 0 of 226,312 non-animal files (Haldane-Anscombe-corrected OR = 23.66; 95% CI 1.45 to 386.0; p = 4.44 × 10^−5^) and 41 of 660,127 versus 0 of 226,312 non-animal files (Haldane-Anscombe-corrected OR = 28.46; 95% CI 1.75 to 461.8; p = 5.64 × 10^−6^) matched occurrences, respectively. These odds ratio’s are in line with other non-alkamines steroidal conjugates such as cholesteryl-linoleate(*38*) that have MS/MS matches in 624 of 660,127 animal-associated data files versus 30 of 226,312 non-animal files (OR = 7.13; 95% CI 4.9 to 5.1; Fisher’s exact test, p = < 1 × 10^− 26^) and lithocholyl amidates(*39*), such as lithocholyl-phenylalanine, that have MS/MS matches in 60 of 660,127 animal-associated files versus 0 of 226,312 non-animal files (Haldane-Anscombe-corrected OR = 3.74; 95% CI 2.5 to 670.9; Fisher’s exact test, p = 8.7 × 10^− 3^). Although this does not take account of addition of feces-based fertilizers to plants or other reasons for unanticipated matches, the animal-restricted prevalence of these scaffolds is consistent with a chemoevolutionary origin, as cholesterol-derived cores are confined to the animal taxonomic branch, emerging only with the advent of animals and their associated gut microbial metabolism(*40*) (**Fig. 4E**). The MS/MS of serine pentadecanal alkamine was detected in 81 of 104,265 plant-associated data files versus 4 of 782,174 non-plant files (OR = 152.03; 95% CI 55.8 to 414.4; p = 6.16 × 10^−70^), and the MS/MS for proline citral in 18 of 104,265 plant-associated files versus 3 of 782,174 non-plant files (OR = 44.98; 95% CI 13.1 to 154.5; p = 1.73 × 10^−14^), with both showing a strong association with plant datasets, consistent with pentadecanal as a plant cuticular wax aldehyde(*41*) and citral as a characteristic citrus-derived monoterpene(*42*). Conversely, MS/MS of the glycine glyceraldehyde and alanine indole pyruvic acid alkamines were detected broadly across different taxonomic groups, reflecting the chemoevolutionary antiquity of their precursors, with glyceraldehyde serving as a conserved glycolytic intermediate and indole pyruvic acid as a tryptophan transamination product, both distributed broadly among bacteria, plants, and animals.

To obtain additional support for these structural hypotheses, biological samples and comparable matrices from humans, lizards, felines and food extracts that had MS/MS matches to the alkamine library were experimentally analyzed by LC-MS/MS alongside authentic synthetic standards, matching MS/MS spectra, retention times, and ion mobility drift times for some specific cases once such instruments became available to our labs. In total, 27 bile acid and steroidal alkamines were supported to level 1 by MS/MS and retention time matching, including 14 of these further supported by ion mobility drift time, detected in cheetah and other large cat fecal extracts, human fecal samples, and cultured microbial communities (**Fig. 4B,F; Fig. S4**,**S5**).

To investigate the microbial origins of these metabolites, a panel of human gut bacterial isolates and defined microbial communities (**Table S17)** that were cultured with and without bile acid supplementation and monitored by LC-MS. Phenylalanine-3-oxo-lithocholic acid production was strictly bile acid-dependent and observed only after 72 h, consistent with an enzymatic biotransformation. Among isolates tested, *B. caccae, B. luhongzhouii, C. innocuum, E. ramosum*, and the microbial community showed the strongest production, *P. distasonis* showed a weaker response, while *B. uniformis* and *P. vulgatus* showed no detectable production (**Fig. 4G**). Concentrations of representative compounds, including phenylalanine-lithocholic acid, phenylalanine-deoxycholic acid, and N-acetylcadaverine-lithocholic acid alkamines, ranged from 1.9 to 14.2 μM in animal fecal samples (**Table S6**). Although the goal of this work is to show that there are entirely overlooked classes of metabolites that were detected but not yet annotated, selected compounds were screened in a general cell-based G protein–coupled receptors (GPCRs) and nuclear receptor-based assay(*43*). The *S*-enantiomer of the alkamine derived from *N*-acetylcadaverine and 3-oxo-lithocholic acid produced the strongest response. (**Fig. 4H,I**). Future work will be needed to identify the specific receptors.

Together, these findings indicate that alkamine formation is not determined solely by the presence of a ketone functional group but instead reflects selective scaffold-dependent transformations. Reductive amination products with steroidal cores are exclusively observed so far at the C3 position of the tetracyclic steroid ring system, with no evidence for modification of side-chain carbonyls or non-C3 ketones.

### Alkamine formation via reductive amination represents a common yet previously undescribed pathway of drug metabolites

Among the drug alkamines detected in public datasets, MS/MS matches to 16 carbonyl-derived 5-ASA alkamines were identified in human samples. Although *N*-acylation products of 5-ASA have been reported(*44, 45*), alkamine derivatives have not. 5-ASA and its prodrugs, sulfasalazine, olsalazine, and balsalazide, are first-line maintenance therapies for inflammatory bowel disease (IBD), particularly ulcerative colitis, and despite this being controversial, they are also prescribed in Crohn’s disease. Azo-linked prodrugs rely on microbial azoreductase activity to release 5-ASA in the colon. Sulfasalazine is additionally used in rheumatoid arthritis (RA), where the sulfapyridine moiety is considered the primary active component(*46, 47*).

Because MS/MS alone cannot always resolve structural isomers, we assessed whether endogenous compounds such as 3-hydroxyanthranilic acid (3-HAA), a tryptophan-kynurenine-NAD+ pathway metabolite(*48*), could account for observed spectral matches. Two independent lines of evidence argue against this. First, 97% of 5-ASA alkamine MS/MS matches occurred in IBD/RA datasets: 50 of 2,193 IBD/RA files contained matches versus only 8 of 503,240 unknown IBD status files (OR = 1,467.7; 95% CI 695 to 3,099; Fisher’s exact test, p = 7.8 × 10^−110^). Restricting to IBD/RA versus healthy individuals within IBD studies, a Haldane-Anscombe-corrected analysis supports strong enrichment (OR = 790; 95% CI 310 to 2,000; p = 3 × 10^−56^), with zero matches in 8,559 LC-MS/MS files from samples originating from healthy individuals. This disease-specific distribution mirrors known 5-ASA prescribing patterns precisely (**Fig. 5A-F**). Second, multiplexed synthetic 3-HAA alkamine standards compared directly against 5-ASA alkamines and human fecal samples by MS/MS, retention time, and ion mobility showed that biological features matched exclusively to 5-ASA alkamines in all cases (**Fig. 5K; Fig. S4**). Nine 5-ASA alkamines achieved level 1 identification by combined MS/MS and retention time matching in rheumatoid arthritis fecal samples, with pyruvate and ribose conjugates quantified at 43.6 to 93.6 μM (**Table S7**), consistent with known 5-ASA exposure in those people from which the samples originated from (**Fig. 5B-F**).

**Fig. 5.**
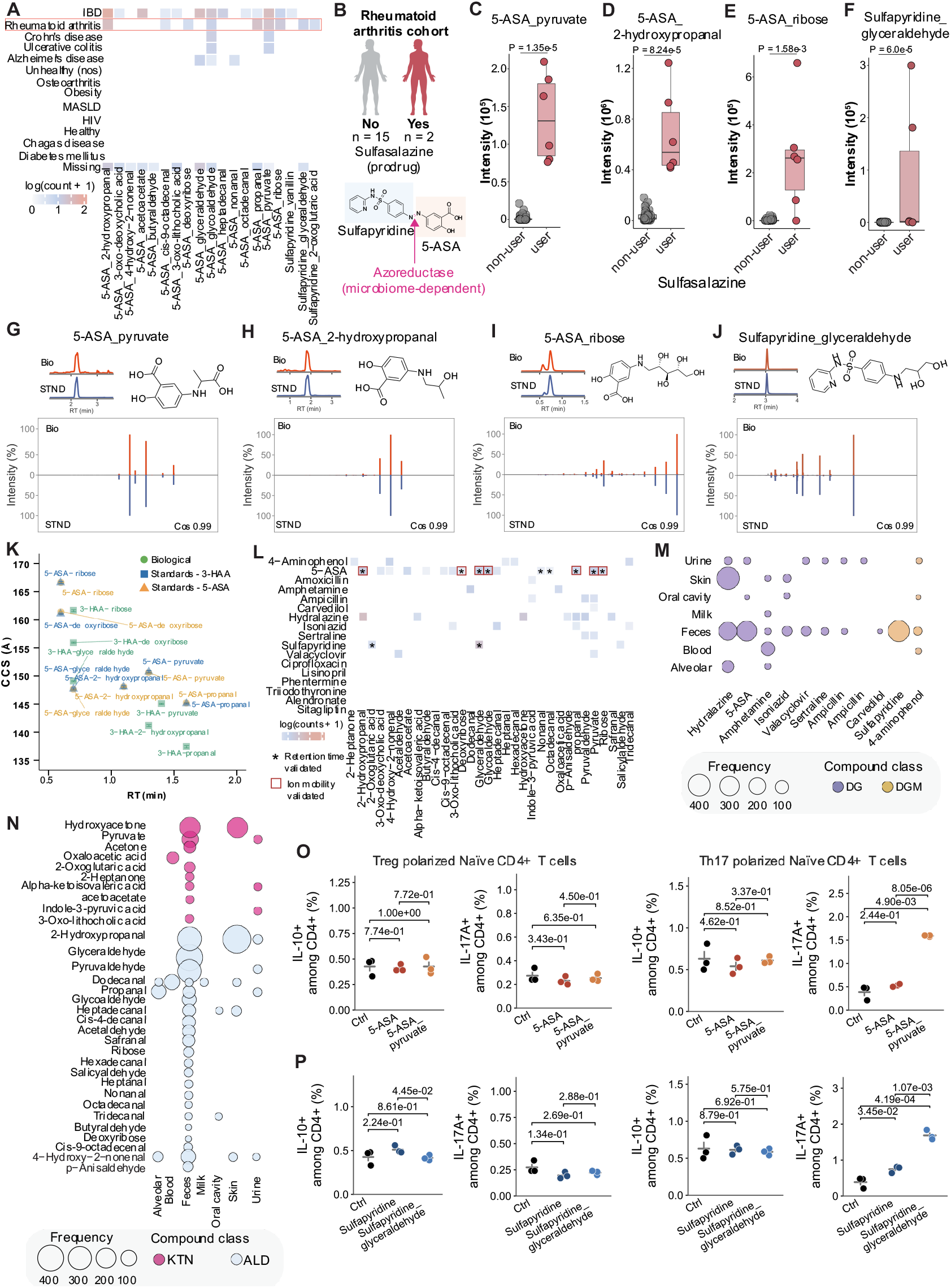
Comparison to standards to support their identification and generalization of alkamine formation as a drug metabolic pathway. **(A)** Abundance, prevalence, and sample-type distributions of 5-ASA and sulfapyridine–derived alkamines across public metabolomics datasets, consistent with known 5-ASA and sulfapyridine exposure in Rheumatoid arthritis(*49*) and IBD in both Crohn’s disease and Ulcerative colitis(*50*). **(B)** Rheumatoid arthritis cohorts with 2/17 individuals reported to be exposed to 5-ASA and sulfapyridine through sulfasalazine drug, each volunteer has 3 biological samples as part of the study that were obtained multiple weeks apart. (**C-F)** Peak areas extracted of 5-ASA and sulfapyridine alkamine conjugates, comparing users of the sulfasalazine colored pink and non-users colored grey for a Rheumatoid arthritis study. **(G-J)** Chromatographic comigration and MS/MS comparison of synthetic 5-ASA and sulfapyridine alkamine standards (STND) and fecal samples of Rheumatoid arthritis human cohorts (Bio), showing exclusive matching to 5-ASA–derived alkamines across retention time and fragmentation. **(K)** Drift time comigration of synthetic alkamine standards and fecal samples of Rheumatoid arthritis human cohorts. **(L)** Heatmap summarizing 148 drug-derived alkamines, organized by amine (y-axis) and carbonyl precursor (x-axis), with a representative 5-ASA and sulfapyridine alkamine conjugates validated by chromatographic coelution represented by a star and drift time represented by a red square. **(M-N)** Bubble plot of drug-derived alkamine occurrence across human body parts and fluid in public metabolomics data, with **M** showing the drugs (DG) and drug metabolite (DGM) part, and **N** showing the ketones (KTN) and aldehydes (ALD) that are being conjugated. **(O)** Naïve CD4⁺ T cells were polarized under Th17 or Treg conditions and treated with 50 µM 5-ASA and 5-ASA_pyruvate, and compared to vehicle control (DMSO, Contr). **(P)** Naïve CD4⁺ T cells polarized under Th17 or Treg conditions were treated with 50 µM sulfapyridine and sulfapyridine_glyceraldehyde, and compared to vehicle control (DMSO, Contr).

In the second phase, we wanted to establish whether other drugs and drug metabolites could undergo this transformation. To assess whether reductive amination represents a broader route of xenobiotic biotransformation, we generated an expanded multiplexed MS/MS library covering 17 amine-containing drugs and metabolites, including 4-aminophenol, sulfapyridine, ampicillin, amoxicillin, amphetamine, valacyclovir, hydralazine, sertraline, and carvedilol, totaling 1,360 putative drug alkamine reference spectra. MS/MS matches to public data were observed for 11 of 17 drugs evaluated, yielding 148 unique drug derived alkamines (**Fig. 5L**). No matches were detected for alkamines of lisinopril, ciprofloxacin, triiodothyronine, alendronate, phentermine, or sitagliptin even though the parent drug was detected in most cases, indicating that alkamine formation is selective and drug-specific rather than universal. Matches were most frequent in feces, followed by urine, skin, and blood (**Fig. 5M,N**).

Additional evidence supporting level 1 identification was obtained for 12 drug-derived alkamines by MS/MS and retention time matching in fecal samples from rheumatoid arthritis patients, with six 5-ASA alkamines further supported by ion mobility drift times that are distinct from corresponding 3-HAA alkamines (**Fig. 5K,L,M; Fig. S6**). Sulfapyridine conjugates to glyceraldehyde and alpha-ketoglutarate were quantified at 3.7 to 11.4 μM in rheumatoid arthritis fecal samples (**Table S5**).

Alkamine modification of both 5-ASA and sulfapyridine altered their immunomodulatory activities on CD4+ T cells *in vitro*. Notably, the alkamine-modified forms (5-ASA pyruvate and sulfapyridine glyceraldehyde) significantly increased IL-17A expression among CD4+ T cells under Th17 polarizing conditions compared to control, whereas the unmodified parent drugs did not produce this effect. This is notable given the pathogenic role of IL-17A in RA and the established Th17-suppressing mechanism of the parent compounds. This suggests that alkamine modification introduces or unmasks molecular interactions that promote rather than suppress Th17 responses. No differences were observed in IL-10 expression under any condition, nor in either cytokine under Treg polarizing conditions, indicating that the effect of alkamine modification may be selectively restricted to IL-17A production in Th17 polarized cells. Together these findings suggest that alkamine drug metabolites may qualitatively alter and in some cases reverse the immunomodulatory properties of the parent compounds, with potential implications for Th17-driven inflammatory diseases such as RA (**Fig. 5O,P, Fig. S9, Table S13**).

Collectively, these findings establish reductive amination as a systematically overlooked pathway of drug metabolism. Unlike canonical phase I and phase II modifications such as acetylation, sulfation, and glucuronidation, which introduce predictable mass shifts from defined enzymatic reactions, reductive amination generates metabolites through combinatorial pairing of amine and carbonyl precursors, producing a structurally diverse and, until now, largely invisible metabolite space. This combinatorial logic likely explains why alkamines have evaded detection and suggests that current drug metabolism frameworks substantially underestimate the chemical diversity of drug-derived metabolites.

## Discussion

Alkamines represent a previously unknown molecular class of the metabolome, now made accessible through the combination of multiplexed synthesis and pan-repository spectral matching. Just as biomedical research has concentrated on a minority of well-studied genes while the functions of most remain uncharacterized, metabolomics has operated under an analogous blind spot: we have measured what we already knew how to find. Their detection across species, organs, and molecular classes demonstrates that reductive amination is a widespread transformation whose products have been systematically absent from metabolomics annotations due to the lack of reference libraries rather than their absence from biology. The picture presented here is necessarily partial. The current library spans 96 aldehydes and ketones and 110 amines, a fraction of the carbonyl and amine-containing molecules present in biology, including endogenous metabolites, microbial products, and drugs. Expanding these libraries will likely reveal many additional alkamines beyond those reported here.

This combinatorial and sample-specific nature of reductive amination is the most likely reason this form of drug metabolism has not been documented before: not everyone makes the same amines, aldehydes, and ketones at the same time, as this depends on fasting state, diet, and microbiome composition, and this observation provided plausible explanations why people respond differently to the same medications and diet (**Fig. S10**). Current drug metabolism frameworks may therefore substantially underestimate the chemical diversity of drug-derived metabolites *in vivo*, with poorly understood consequences for pharmacology, toxicology, and drug-drug interactions. The discovery of alkamines across diverse species, organs, and molecular classes reveals that the annotated metabolome systematically excludes an entire class of biologically produced molecules, not because they are rare or analytically inaccessible, but because reference libraries were never built for them. The field has been looking under the streetlight: reporting only to what can already be annotated, while leaving the rest unseen. Genomic and metagenomic surveys have identified imine reductases across all domains of life, including in humans, establishing that the enzymatic capacity for reductive amination is broadly distributed in biology. Yet because the synthetic biology and pharmaceutical communities developed these enzyme inventories solely to manufacture chiral amines, no one asked what this chemistry produces endogenously. By revealing, in 2026, more than 80 years after the first metabolic pathways were drawn, a class of metabolites that is both chemically widespread and systematically absent from existing reference libraries, this work exposes a structural blind spot in how the field detects, identifies, and interprets small molecules in biological systems. The gap is not one of analytical sensitivity but of chemical imagination: the metabolome has always contained these molecules, but the frameworks for finding them did not.

The broader significance extends beyond alkamines themselves. Reductive amination is likely one of many transformations seen in biological systems whose products remain undiscovered by standard workflows for the same reason: the absence of reference standards rather than the absence of reactivity and biology. This suggests that the dark matter of the metabolome is not simply uncharacterized, but is in part the predictable output of chemistry that has not yet been systematically anticipated. Just as Richardson and colleagues showed that understudied genes are not absent from experimental data but are abandoned before results are reported(*51*), the alkamines reported here were present in the data all along, detectable in over 1.7 billion publicly available spectra, yet invisible without the right reference framework. Multiplexed synthesis paired with pan-repository spectral matching offers a generalizable strategy for illuminating these regions, one that can be extended to other reactive chemistries, expanded amine and carbonyl libraries, and additional biological contexts.

When metabolite inventories are incomplete, biological interpretation becomes fragile, akin to reading a book with 93% of the words missing. In fundamental biology and medical research, incomplete chemical maps obscure connections between host, microbiome, diet, and environment and limit the interpretation of disease-associated metabolites. Many biologically informative molecules arise from medications, dietary preferences, microbiome composition, or environmental exposures and therefore appear only in subsets of individuals within a cohort. Yet although infrequent in a single study, many are common across populations and datasets, making their annotation essential for interpreting metabolomics at scale. For drug metabolism in particular, the consequences are immediate. A pharmacology built on incomplete inventories of metabolic products risks misattributing biological effects, overlooking toxic species, and failing to anticipate interactions arising from combinatorial biotransformation products. Closing this gap will require not only larger libraries, but a shift in the conceptual scaffolding of metabolomics toward frameworks that treat unannotated chemical space not as an intractable unknown, but as a chemically predictable territory that can be mapped using data science workflows that leverage raw data to systematically build reference libraries, as demonstrated here.

## Supporting information

Supplementary Materials including methods and figures

## Author contributions

Conceptualization, JA and PCD; Methodology, JA; LC-MS/MS data collection, JA, VCL and AP; Data analysis, SX, VCL, HM-R, MRN, YEA, JS, HG; Chemical synthesis, SRP, AP, SG, ZH; immune T-cell screening; MCT, HC, Nuclear receptor screening; RND, CKA; Microbial culturing, VCL; Resources, KEK, HNZ, IM, SKY, JC, HMR, IS, RB, AMCR, CEW, CLW, JC, NR, KD, WDGN, LAB, AJ, IAM, NB, GP, LRH, RK, MW, MSC, MG, and PCD; Library creation, JA, JZ, PR and MW; Writing original draft, JA and PCD; Writing, review & editing, all authors; Supervision and funding acquisition, PCD and DS.

## Statistics

Enrichment of 5-ASA alkamine MS/MS matches in disease-associated datasets was assessed using two-sided Fisher’s exact tests applied to contingency tables. Effect sizes were quantified as odds ratios (ORs) with corresponding 95% confidence intervals (CIs). In cases involving small or zero cell counts, a Haldane–Anscombe correction was applied prior to OR estimation to reduce bias and ensure numerical stability. Comparisons between USER and Non-USER groups were performed using the Wilcoxon rank-sum test for continuous variables and Fisher’s two-sided exact tests for categorical variables, including analyses of sulfapyridine_glyceraldehyde detection.

## Data availability

The data is publicly available on GNPS/MassIVE: Untargeted metabolomics reanalysis: MSV000084556 (rheumatoid arthritis); Unprocessed data to alkamine library: MSV000095820 and the library itself is available is part of the GNPS2 reference library https://library.gnps2.org/; Comigration of synthetic alkamine and biological samples: MSV000095820, MSV000099877, MSV000099783, MSV000100346, MSV000099653, MSV000100140, MSV000100138, MSV000099770, MSV000100141, MSV000100138, MSV000099967, MSV000099978, MSV000100441, MSV000100287, MSV000100340, MSV000100197, MSV000100532, MSV000100534, MSV000101107; Ion mobility of drug and steroids alkamines; Quantification of drug and steroids alkamines: MSV000101093. NMR data are available on Zenodo (NMR data for Alkamine).

## Code availability

The synthesized alkamine library was queried against public metabolomics repositories using fast MASST (FASST) (https://github.com/robinschmid/microbe_masst). Structural overlap with public spectral libraries was assessed, alongside species-level categorization, feature counts, and validation of FASST-derived matches https://github.com/helenamrusso/cosine_calculation_FASST_search). The distribution of annotated alkamines was further examined across human body fluids and organs, as well as within microbial culture datasets. Additional analysis pipelines are available at https://github.com/Philipbear/alkamine_analysis. Comparative analyses of sulfasalazine users and non-users in a rheumatoid arthritis cohort, https://github.com/VCLamoureux/Alkamines.git, filtering bile acid alkamines peaks from FASST data https://github.com/jagongo94/Diagnostic-bile-acid-MS-MS-spectrum-filter

## Use of AI and Software with AI for the research

In the Dorrestein Lab, the use of AI and AI-enabled tools, including generative AI, large language models (LLMs), and software that leverages these technologies, both free and commercial, is encouraged across all aspects of research. These tools support a wide range of activities, including literature searches, data analysis, scripting, coding, text editing, and figure concept development. All figures and analyses are original. To ensure transparency and reproducibility, all raw data, derived data tables and code used in this research are made accessible and linked with this manuscript.

