## Supplementary Materials including methods and figures for "Alkamines reveal a hidden layer of steroid and drug metabolism"

The PDF file includes:

Materials and Methods
Supplementary Text
Figs. S1 to S21
Table S1 to S17 are provided as an Excel file
References

### Materials and Methods

Multiplexed synthesis of alkamines

96 different Aldehydes or ketones containing compounds were weighed and dissolved in methanol (MeOH) to make a stock concentration of 10mM. Subsequently, 400uL of 100 µM was prepared for each aldehyde or ketone via serial dilution and separately added into a well in a deep 96 well plate. A total of 110 different amines were used for the reaction. Briefly, 400uL of a mixture of amines, 20 in number for each batch of the mixture, was added into each well containing the aldehyde or ketone to make a total molar equivalence of 1.5:1 for aldehyde or ketones/amines. 80 uL of acetic acid was added to each reaction to catalyze the imine formation. The reaction was let to sit at room temperature for 30 minutes followed by addition of 3 equivalents of sodium cyanoborohydride. The reaction was further stirred for 30 minutes and then quenched with 100uL of saturated ammonium bicarbonate buffer. The mixture was then concentrated using a vacuum centrifuge concentrator (room temperature; ~3 hours) and stored at -80 °C. List of aldehydes, ketones and amines used for the reactions is found (**Table S9**).

LC-MS/MS analysis for multiplex synthesis of alkamines

The reaction mixture was reconstituted into 100% MeOH containing 0.1% formic acid and transferred into a 96 shallow well plate for analysis. A volume of 5uL was injected into a Vanquish ultra-high-performance liquid chromatography (UHPLC) system coupled to a Q Exactive quadrupole Orbitrap mass spectrometer (Thermo Fisher Scientific, Waltham, MA). For the polar synthesized alkamines, an ACQUITY PRM Gly BEH amide HILIC column (2.1 × 100mm, 1.7 µm particle size, 130 A pore size; Waters) was used at 40 °C column temperature to analyze them. A flow rate of 0.4 mL/min used for a 12-minute total run time. The mobile phase A and B were 0.1% formic acid/water and 0.1% formic acid/acetonitrile respectively, gradients for the run are as follow: 0-2 min 5 % A, 2-7 min 5-40 % A, 7-10 min 40-98% A, and 10-12 min 5% A for re-equilibration. For the nonpolar synthesized alkamines, a Kinetex non-polar C18 column (2.1 × 100 mm, 2.6 µm particle size, 100 A pore size; Phenomenex, Torrance) was used together with a SecurityGuard C18 column (2.1 mm ID) at 40 °C column temperature. A flow rate of 0.5 mL/min used for a 12-minute total run time. The mobile phase

A and B were 0.1% formic acid/water and 0.1% formic acid/acetonitrile respectively, gradients for the run are as follow: 0-1 min 5 % B, 1-7 min 5-40 % B, 7-10 min 40-98% B, and 10-12 min 5% B for re-equilibration. A positive mode electrospray ionization was used analyze all samples with the following parameters: sheath gas flow, 53 AU; auxiliary gas flow, 14 AU; sweep gas flow, 3 AU; auxiliary gas temperature, 400 °C; spray voltage, 3.5 kV; inlet capillary temperature, 269 °C; S-lens level, 50 V. MS1 scan was performed at m/z 100-1500 with the following parameters: resolution, 17,500 at m/z 200; maximum ion injection time, 100 ms; automatic gain control (AGC) target, 1E6. Up to 5 MS/MS spectra per MS1 scan were recorded under the data-dependent mode (dd-MS2) with the following parameters: resolution, 17,500 at m/z 200; maximum ion injection time, 150 ms; AGC target, 2.0E5; inclusion set on for list of 200 target m/z, MS/MS precursor isolation window, m/z 1; isolation offset, m/z 0.5; stepped normalized collision energy (NCE) of 25, 40 and 60%; minimum AGC for MS/MS spectrum, 5E4; apex trigger, 2 to 5 s; dynamic precursor exclusion, 10 s.

MS/MS spectra library generation for synthesized alkamines

Creation of SMILES for the expected products: A total of 13110 SMILES were generated for expected products using smiles of the starting materials and a smart reaction. The SMILES strings were created using a script that can be accessed through this link https://autosmiles.streamlit.app/rxnSMILES. All SMILES for the expected product can be found (**Table S10**).

MS/MS extraction: the Thermo raw files containing MS/MS spectra were converted to mzML files using MSconvert (ProteoWizard) peak picking. An annotation table containing name, SMILES and molecular formula for targeted alkamines synthesized products was created and used together with the mzML file to generate a spectra library using our unpublished but publicly accessible workflow "reverse metabolomics create library" in GNPS2, GNPS2 - Analysis Hub. Briefly minimum feature height was set to 1.5e5, ms2 explanation cutoff 0.9 out of 1. The tsv output from the workflow containing extracted MS2 for the synthesized compounds was used to generate the mgf spectra library for alkamines. A total of 18,031 MS/MS spectra summarized to 8,475 unique compounds were recovered from the synthesis. The links for the synthesized alkamine libraries can be found here: https://gnps.ucsd.edu/ProteoSAFe/gnpslibrary.jsp?library=GNPS-ALKYLAMINES-BILE-ACIDS-LIBRARY and https://gnps.ucsd.edu/ProteoSAFe/gnpslibrary.jsp?library=GNPS-ALKYLAMINES-LIBRARY

Repository MS/MS search of synthesized alkamines

Reference MS/MS spectra from the synthesized alkamine library were searched against public metabolomics repositories using fast MASST (FASST)(*1*). FASST compares query spectra against LC-MS/MS data across public repositories and identifies matching spectra based on spectral similarity. In this study, searches were performed in a batch using microbe_masst in Python. Matches were required to have a minimum cosine score of 0.8, at least four matched fragment ions, an MS1 tolerance of 0.02 Da, and an MS/MS tolerance of 0.02 Da.

FASST returns matching MS/MS scans along with their associated dataset and file identifiers. When available, domain-level MASST annotations were used to classify matching files as microbe-, plant-, or food-associated(*2*). To evaluate phenotype associations, MASST results were merged with the Pan-ReDU metadata table. Disease-related annotations were extracted from the DOIDCommonName and HealthStatus fields. Body site annotations were obtained from UBERONBodyPartName, and taxonomic annotations were obtained from NCBITaxonomy.

Filtering of the Pan-repository results

*Cosine recalculation of Pan-repository results to reevaluate pre-filtered spectra:* FASST searches rely on pre-filtered spectra, and to ensure higher quality matches, the cosine scores of unfiltered spectra were also calculated. Raw cosine scores were recalculated for each scan retrieved from the pan-repository against the synthesized alkamines. Here, MS/MS spectra were downloaded and mirror-matched against the reference spectra to compute raw cosine values using the Python code similar to previous study(*3*). The resulting raw cosine scores were filtered using a minimum threshold of 0.8 and at least four matching peaks.

*MassQL for capturing bile acids alkamine using diagnostic peaks:* Bile acid alkamine matches from the pan-repository results were checked based on diagnostic fragment ions at *m/z* 359.294, 357.278, and 355.260, corresponding to 3-position–alkylated lithocholic acid, deoxycholic acid, and cholic acid, respectively(https://github.com/jagongo94/Diagnostic-bile-acid-MS-MS-spectrum-filter). Briefly, a ratio of the diagnostic peaks to the common peaks that exist between the alkamine and the amidate (341.2831, 323.2722 for lithocholic acid, 339.2667, 321.2561 for deoxycholic acid and 337.2510, 319.2405 for cholic acid) and filter for true positive if the ratio ≥ 1 This additional filtering step was applied to distinguish bile acid alkamines from bile acid amides, which are structural isomers that produce overlapping fragment ions. For bile acid alkamines, a lower cosine score threshold (≥0.6) was accepted only when diagnostic fragment peaks were present.

Reanalysis of public dataset from GNPS/MassIVE

Publicly available stool untargeted metabolomics data from a rheumatoid arthritis cohort (MSV000084556) was reanalysed to evaluate the presence of the 5-ASA derivatives from our repository-scale analyses as described previously(*4*). Briefly, the data (.mzML files) were downloaded from GNPS2/MassIVE, and the features were extracted using MZmine 4 (version 4.5.0). An mgf containing the spectra (list of *m/z* and matching intensities) and csv files (containing the relative peaks areas) were obtained from the batch processing workflow, followed by a feature-based molecular networking analysis (FBMN) on GNPS2. The FBMN parameters were set to minimum cosine of 0.7, matching peaks 4, and precursors and fragment tolerance of 0.02 and run against the alkamines synthesized library available here: https://gnps2.org/status?task=d7c5882a01954ed28294f340dd855ee2. The quant table as well as the annotation table was imported in RStudio for downstream analyses. For the bacterial culture, the same process was used as above and the culturing details can be found elsewhere(*5*). Briefly, the data was downloaded from GNPS2/Massive (MSV000098955), processed with MZmine for feature detection. Tables generated by MZmine were imported into RStudio for downstream analysis.

LC-MS/MS data acquisition for co-migration of synthetic standards and biological samples

Comigration of synthesized alkamine standards and fecal samples from humans was performed for level identification. Briefly, for each well of the 96-well plate containing the alkamine of interest, 10 µL of the synthesized standard in 100% methanol (MeOH) was diluted 100-fold and analyzed by LC–MS/MS. Similarly, 100 µL of extracts from human fecal; Diet_009, Diet_007, RA_RDT_P4-A6 and RDT-P3_F10, obtained from a rheumatoid arthritis cohort (UC San Diego IRB# 161474, and IRB# 191900), HNRC_P3_E10 and HNRC_P4_E2 obtained from a Human Immunodeficiency Virus cohort (UC San Diego IRB# 172092), and P1-E11-311_2, P2-E12-334_2 obtained from inflammatory bowel disease cohort (UC San Diego IRB# 800805) study were dried down, reconstituted in 100% MeOH, and analyzed by LC-MS/MS under the same chromatographic and MS/MS conditions described below; a volume of 5 µL was injected into a Vanquish ultra-high-performance liquid chromatography (UHPLC) system coupled to a Q Exactive quadrupole Orbitrap mass spectrometer (Thermo Fisher Scientific, Waltham, MA). A Kinetex non-polar C18 column (2.1 × 100mm, 2.6 µm particle size, 100 A pore size; Phenomenex, Torrance) was used together with a Security Guard C18 column (2.1 mm ID) at 40 °C column temperature. A flow rate of 0.5 mL/min used for a 12-minute total run time. The mobile phase A and B were 0.1% formic acid/water and 0.1% formic acid/acetonitrile respectively. The gradients for the run are as follows: 0-1 min 2 % B, 1-2.5 min 2-20 % B, 2.5-4 min 20% B, 4-6 min 20-98% B, 6-10 min 98% B and 10-12 min 2% B for re-equilibration. A positive mode electrospray ionization was used with the following parameters: sheath gas flow, 53 AU; auxiliary gas flow, 14 AU; sweep gas flow, 3 AU; auxiliary gas temperature, 400 °C; spray voltage, 3.5 kV; inlet capillary temperature, 269 °C; S-lens level, 50 V. MS1 scan was performed at m/z 100-500 with the following parameters: resolution, 17,500 at m/z 200; maximum ion injection time, 100 ms; automatic gain control (AGC) target, 1E6. Parallel reaction monitoring (PRM) was used for MS/MS with the following parameters: resolution, 17,500 at *m*/*z* 200; maximum ion injection time, 100 ms; AGC target, 2.0E5; MS/MS precursor isolation window, *m*/*z* 1; isolation offset, *m*/*z* 0.0; stepped normalized collision energy (NCE) of 20, 30 and 40% for the target list of alkamines.

LC-IM-MS/MS data acquisition for comigration of synthetic standards and biological samples. Some of the alkamines were further validated by comigration with alkamine standards, and their presence in human fecal extracts was additionally confirmed using ion mobility spectrometry. Analyses were performed on an Agilent liquid chromatography system coupled to a hybrid trapped ion mobility–quadrupole time-of-flight mass spectrometer (timsTOF Pro II). Chromatographic separation was achieved using a non-polar Kinetex C18 column (2.1 × 100 mm, 2.6 µm particle size, 100 Å pore size; Phenomenex) equipped with a matching C18 guard cartridge (2.1 mm internal diameter). The mobile phases consisted of solvent A (water containing 0.1% formic acid) and solvent B (acetonitrile containing 0.1% formic acid). The column oven temperature was maintained at 40 °C. Samples (5 µL) were injected and eluted at a flow rate of 0.5 mL/min using the following gradient program: 5% B from 0–0.5 min; 25% B from 0.5–1.1 min; 40% B from 1.1–7.5 min; 99% B from 7.5–8.5 min, held at 99% B until 10 min; followed by re-equilibration to 5% B from 10–10.1 min and maintained at 5% B until 12 min. The QTOF mass spectrometer was pre-calibrated in positive ion mode using sodium formate, achieving a mass accuracy with a standard deviation below 0.5 ppm. Mass spectra were acquired over an m/z range of 20–1300 using parallel accumulation–serial fragmentation (PASEF) mode. Instrument parameters included an end plate offset of 500 V, capillary voltage of 4500 V, nebulizer pressure of 2.2 bar, dry gas flow of 10 L/min, and a dry gas temperature of 200 °C. Low-abundance precursor ions with intensities exceeding 100 counts but below a target threshold of 4000 counts were selected for fragmentation, with an active exclusion window of 0.05 min. Collision-induced dissociation was performed using collision energies of 50 and 60 eV.

Quantification of Steroid and Drug-Conjugated Alkamines in humans and feline fecal Samples

Absolute quantification of steroid alkamines (phenylalanine_3-oxo-lithocholic acid, phenylalanine_3-oxo-deoxylithocholic acid, and N-acetylcadaverine-lithocholic acid) in human and feline fecal samples, as well as drug-conjugated alkamines (5-aminosalicylic acid_pyruvate, 5-aminosalicylic acid_ribose, sulfapyridine_glyceraldehyde, and sulfapyridine_alpha ketoglutaric acid) in human fecal samples, was performed using matrix-matched calibration curves.

Individual analytes were weighed using an OHAUS™ 30100600/EMD microscale analytical balance (Fisher Scientific, USA) with a precision of 0.0001 g and dissolved in 1 mL methanol/water (50:50) to prepare 1 mM stock solutions. A combined working solution (100 µM) was subsequently prepared, followed by serial dilution to generate calibration standards at 0.01 µM, 0.1 µM, 0.2 µM, 0.4 µM, 1.0 µM, and 2.0 µM. Pooled fecal samples were spiked into each level to produce matrix-matched calibrators. A non-isotopic internal standard (sulfadimethoxine) was prepared similarly and added to all samples at a final concentration of 1 µM. LC–MS data acquisition was performed as described above. Peak areas were extracted from raw Thermo files using Skyline (version 21.2.0.425)(*6*). Peak area ratios were calculated by dividing the analyte signal by that of the internal standard(*7*).

Calibration curves were generated using these ratios and corresponding concentrations. A weighted second-degree polynomial regression model (inverse response weighting) was applied, particularly improving fit at lower concentrations. Quadratic equations derived from these models were used for quantification. The limits of detection (LOD) and quantification (LOQ) were calculated as LOD = 3.3 ($\sigma$/S) and LOQ = 10 ($\sigma$/S), where $\sigma$ represents the standard deviation of the y-intercept and S the slope of the calibration curve. Precision was assessed as the coefficient of variation (CV), and accuracy was determined by comparing measured concentrations to theoretical values (n = 3 injections). For steroid alkamines, quantification was performed in human samples (HNRC_P4E2) and a feline sample (AL-3-Cat, African lion). For drug-conjugated alkamines, quantification was performed in human fecal samples (Diet_009 and Diet_007). In all cases, calculated concentrations were adjusted using appropriate dilution factors to reflect physiological levels. Replicate injections ($n \geq 3$ per sample) were used to calculate CV values (**Table S6 and S7**).

Immunomodulatory activity Naïve $CD4^+$ T cell assay for 5-ASA and Sulfapyridne alkamines

Co-culture of bone marrow-derived dendritic cells (BMDCs) and bulk CD4+ T cells were performed previously described(*8*). Briefly, BMDCs were generated from bone marrow progenitor cells isolated from femurs of WT C57BL6/J mice in the presence of 20 ng/ml GM-CSF (Miltenyi) in complete RPMI 1640 (10 % fetal bovine serum, 50 U/mL penicillin, 50 µg/mL streptomycin, 2 mM L-glutamine, 1 mM sodium pyruvate, 1 mM HEPES, non-essential amino acids, and beta-mercaptoethanol). BMDCs co-cultured with splenic CD4+ T cells from Foxp3$^{hCD2}$/IL-10$^{Venus}$ (Keio University, Tokyo Japan)(*9*) at a ratio of 1:10 (DC:CD4+ T cells) and treated with 50μM of selected compounds for T helper 17 polarizing conditions in the presence of anti-mouse CD3 (2 μg/ml, eBiosciences), mouse IL-2 (10 μg/ml, Peprotech), mouse IL-6 (20 ng/mL, Peprotech), mouse IL-1b (20 ng/mL, Peprotech), mouse IL-23 (10 ng/mL, Life Technologies) and human TGF-beta (2 ng/ml, Peprotech). For T regulatory polarizing conditions in the presence of anti-mouse CD3 (0.033μg/mL, eBiosciences), mouse IL-2 (1.331μg/mL, Peprotech), and human TGF-beta (5ng/mL, Peprotech). After 3 days of co-culture, cells were stimulated with a cell activation cocktail with Brefeldin A (2 μL/ml, Biolegend) for 4 hours and stained for 30 min at 4 ºC with either LIVE/DEAD fixable yellow dead stain Kit (Life Technologies), with empirically titrated concentrations of the following antibodies: APC-eFluor780-conjugated anti-mouse CD4 (clone: RM4-5). For intracellular staining, cells were fixed and permeabilized using the Foxp3/Transcription Factor Staining Buffer Set (Life Technologies). We followed the manufacturer's protocol, incorporating previously described modifications for staining cytosolic reporter proteins(*10*). Intracellular staining was performed with PE-Cy7-conjugated anti-mouse IL-17A (clone: eBio17B7, and BrilliantViolet785-conjugated anti-mouse IFNγ (clone: XMG1.2) for 30 minutes. All antibodies were purchased from Thermo Scientific/eBiosciences, and Biolegend. Cell acquisition was performed on Fortessa (BD), and data was analyzed using FlowJo software suite (TreeStar) (**Table S13).**

#### HL-60 Calcium Mobilization cell-based assay for estimation of steroid alkamines inflammatory action

Human promyelocytic leukemia cell line (HL-60 cells) maintained in complete RPMI 1640 (10 % fetal bovine serum, 50 U/mL penicillin, 50 μg/mL streptomycin) were transferred into in 10% charcoal-stripped FBS, phenol red-free RPMI for 72 h as described in previous study(*11*). Cells were then treated with 5 μM Indo-1 AM (AAT Bioquest, Inc), 0.05% pluronic acid and 2.5 mM probenecid in 50:1 HBSS-HEPES (Thermo Fisher Scientific, Waltham, MA) and vortexed for 0.5 h at room temperature. Cells were spun down and washed with 50:1 HBSS-HEPES and resuspended in 50:1 HBSS-HEPES. The resuspension was placed on ice for 5 min followed by incubation for 15 min at 37°C before compound addition in the Flex Station 3 Multimode Plate Reader (Molecular Devices, Sunnyvale, CA) for 150 s at 37°C. Calcium mobilization was determined ratiometrically at $\lambda_{ex}$ 350 nm and $\lambda_{em}$ 405/490 nm, respectively (**Table S14)**

#### Targeted chemical synthesis and NMR Characterization

Commercially available reagents and analytical-grade solvents were used without further purification, and all reactions were monitored by TLC visualized under UV light or with cerium ammonium molybdate stain; $^{1}$H and $^{13}$C NMR spectra were recorded on Bruker AVANCE III 600 MHz fitted with1.7 mm probe and 600 MHz fitted with 5 mm triple-resonance cryoprobe instruments as well as on a JEOL 500 MHz spectrometer (5 mm probe), with chemical shifts reported in ppm relative to the residual solvent signals—$CDCl_3$ ($\delta = 7.26$ ppm for $^{1}$H, $\delta \approx 77.0$ ppm for $^{13}$C), methanol-$d_4$ ($\delta \approx 3.31$ ppm for $^{1}$H, $\delta \approx 49.0$ ppm for $^{13}$C), and DMSO-$d_6$ ($\delta \approx 2.50$ ppm for $^{1}$H, $\delta \approx 40.0$ ppm for $^{13}$C; coupling constants (J) are expressed in Hertz, and multiplicities are denoted as s = singlet, d = doublet, t = triplet, q = quartet, m = multiplet, dd = doublet of doublets. Column chromatographic purifications were performed on a

Teledyne Isco CombiFlash Rf system using pre-packed silica ($SiO_2$) columns with dichloromethane/methanol as the mobile phase, while reverse-phase separations employed pre-packed C18 columns with an acetonitrile/water gradient, optionally containing TFA as a buffer additive.

*Synthesis of 3-oxolithocholic acid*

A solution of lithocholic acid (5.0 g, 13.3 mmol, 1 equiv) in dry dichloromethane (100 mL) was prepared under nitrogen at room temperature. To this solution, silica gel (20 g) was added, followed by pyridinium chlorochromate (6.49 g, 17.3 mmol, 1.3 equiv). The reaction mixture was stirred at ambient temperature overnight. Completion of the oxidation was confirmed by TLC. The reaction mixture was filtered through a Celite® pad, and the filtrate was washed sequentially with 5 % MeOH/DCM. The filtrate was washed with 100ml of 1 M HCl, 100ml of water, and 50ml of brine. The organic layer was dried over anhydrous $Na_2SO_4$, filtered, and concentrated under reduced pressure to give the crude product. Purification was performed by flash column chromatography on silica gel, eluting with a gradient of 0 % to 3 % MeOH in DCM. Fractions containing the desired product were combined and concentrated to afford 3-oxolithocholic acid (3.1 g, 62) %).

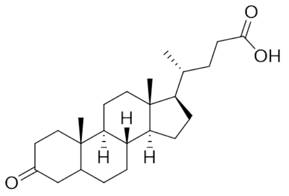


$^{1}$H NMR (500 MHz, DMSO-$d_6$) δ 11.96 (s, 1H), 2.74 (t, $J$ = 14.1 Hz, 1H), 2.36 (td, $J$ = 14.6, 5.4 Hz, 1H), 2.23 (ddd, $J$ = 15.2, 9.7, 5.3 Hz, 1H), 2.15 – 2.07 (m, 1H), 2.01 – 1.90 (m, 1H), 1.88 – 1.76 (m, 2H), 1.76 – 1.63 (m, 2H), 1.62 – 1.50 (m, 2H), 1.45 – 1.35 (m, 3H), 1.33 – 1.03 (m, 14H), 0.96 (s, 3H), 0.88 (d, $J$ = 6.5 Hz, 3H), 0.64 (s, 3H).

*Synthesis of 3-oxo-deoxycholic acid*

A solution of deoxycholic acid (5.00 g, 12.7 mmol, 1 equiv) was placed in a 250 mL round-bottom flask, diluted with anhydrous methanol (50 mL) and p-toluenesulfonic acid monohydrate (242 mg, 0.10 equiv). The mixture was sonicated at ambient temperature for 2 h, after which the solvent was removed under reduced pressure at 40 °C. The residue was dissolved in anhydrous dichloromethane (500 mL) and transferred to a separatory funnel, washed sequentially with 200 mL of saturated aqueous $NaHCO_3$, 200 mL of deionized water, and 100 mL of brine; the organic layer was then dried over anhydrous $Na_2SO_4$, filtered, and concentrated to give the methyl deoxycholate (4.2 g, 84%). Methyl deoxycholate (2.0 g, 4.9mmol, 1.0 equiv) was dissolved in toluene (50 mL) in a round-bottom flask equipped with a Dean–Stark trap; 5.4 g of Fetizon's reagent (2 equiv, 9.8 mmol $Ag_2CO_3$, 50%wt/wt) was added, the suspension was heated to reflux and water formed during the reaction was removed continuously by Dean–Stark distillation; after 16 h at reflux TLC indicated complete consumption of the starting ester, the mixture was cooled to ambient temperature, filtered through a Celite pad and the cake was washed with dichloromethane; the combined filtrate was concentrated under reduced pressure to give the crude ketone, which was purified on silica gel using an ethyl acetate-hexane gradient (0 %-50 % ethyl acetate in hexanes) to afford methyl-3-

oxo-deoxycholate (1.3 g, 65 %). To a solution of methyl-3-oxo-deoxycholate (1.0 g, 1 Eq, 2.47 mmol) in MeOH ( 20 mL) was added NaOH (494.3 mg, 5 Eq, 12.36 mmol) and stirred for 5 h at rt. The reaction was monitored by TLC. Reaction mixture was concentrated to remove the solvent, then diluted with 100ml of water and acidified using dil. HCl, resulted solid was extracted 100 mlx2 of DCM and organic layer was dried over sodium sulfate and concentrated to get the 3-oxo deoxycholic acid (950 mg, 98.4 %) .

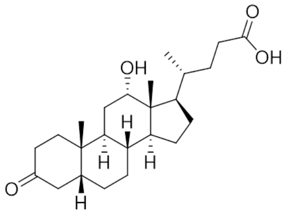

$^{1}$H NMR (600 MHz, Chloroform-*d*) δ 4.05 (t, *J* = 2.8 Hz, 1H), 2.79 – 2.67 (m, 1H), 2.40 (tdd, *J* = 20.1, 12.3, 5.3 Hz, 6H), 2.28 (ddd, *J* = 15.9, 9.5, 6.7 Hz, 3H), 2.16 (dd, *J* = 11.9, 3.2 Hz, 2H), 2.01 (dddd, *J* = 22.7, 14.1, 5.1, 2.9 Hz, 4H), 1.96 – 1.77 (m, 10H), 1.73 – 1.44 (m, 8H), 1.43 – 1.23 (m, 9H), 1.12 (dtq, *J* = 22.5, 11.9, 5.0 Hz, 4H), 1.03 – 0.92 (m, 6H), 0.72 (s, 3H).

*Synthesis of Phenylalanine_3-oxo-lithocholic acid (N-(3-lithocholyl)phenylalanine):*

To a stirred solution of 3-oxolithocholic acid (500 mg, 1 Eq, 1.33 mmol) in methanol (20 mL) was added acetic acid (802 mg, 764 µL, 10 Eq, 13.3 mmol) followed by L-phenylalanine (662 mg, 3 Eq, 4.00 mmol). The reaction mixture was stirred at room temperature for 2 h. Sodium cyanoborohydride (252 mg, 3 Eq, 4.00 mmol) was then added, and the reaction was stirred at room temperature overnight. Reaction progress was monitored by LC–MS. Upon completion, the solvent was removed under reduced pressure to remove methanol and excess acetic acid. The crude residue was diluted with a mixture of acetonitrile/water and purified by reverse-phase flash chromatography (C18 column) using 60–70% acetonitrile/water (0.1% TFA additive) as gradient elution. Fractions containing the desired products were combined and concentrated to afford the following diastereomers:

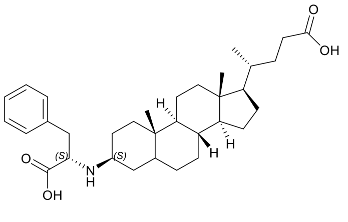

*N*-[(3S)-lithocholyl]phenylalanine (195 mg, 372 µmol, 27.9 %): $^{1}$H NMR (600 MHz, Methanol-$d_4$) δ 7.30 – 7.20 (m, 5H), 4.12 (t, *J* = 7.6 Hz, 1H), 3.45 (s, 1H), 3.20 – 3.07 (m, 1H), 2.22 (ddd, *J* = 15.2, 9.7, 5.2 Hz, 1H), 2.09 (ddd, *J* = 15.5, 9.3, 6.7 Hz, 2H), 1.91 (d, *J* = 12.8 Hz, 1H), 1.82 – 1.74 (m, 3H), 1.74 – 1.66 (m, 2H), 1.63 (d, *J* = 13.4 Hz, 1H), 1.58 (s, 1H), 1.50 (dt, *J* = 9.7, 4.9 Hz, 1H), 1.35 (td, *J* = 19.9, 12.9 Hz, 6H), 1.21 (tdd, *J* = 22.7, 11.0, 7.0 Hz, 9H), 1.14 – 0.92 (m, 8H), 0.89 (s, 3H), 0.84 (d, *J* = 6.5 Hz, 3H), 0.59 (s, 3H). $^{13}$C NMR (151 MHz, Methanol-$d_4$) δ 176.8, 169.3, 134.2, 129.0, 128.8, 127.6, 60.0, 56.2, 56.0, 55.8, 42.5, 40.0, 39.9, 36.2, 35.7, 35.4, 35.3, 34.5, 30.9, 30.6, 29.6, 27.8, 26.6, 26.0, 25.6, 23.8, 22.4, 22.0, 20.6, 17.3, 11.0.

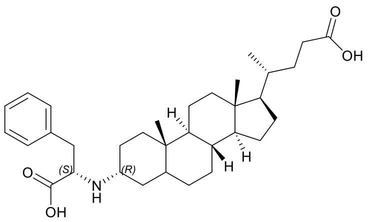

*N*-[(3R)-lithocholyl]phenylalanine (180 mg, 344 μmol, 25.7 %) $^{1}$H NMR (600 MHz, Methanol-$d_4$) δ 7.39 – 7.28 (m, 5H), 4.35 (t, $J$ = 6.8 Hz, 1H), 3.29 – 3.18 (m, 2H), 3.14 (td, J = 12.3, 5.9 Hz, 1H), 2.32 (ddd, $J$ = 15.1, 9.7, 5.2 Hz, 1H), 2.20 (dt, $J$ = 15.5, 8.2 Hz, 1H), 2.06 – 1.83 (m, 8H), 1.83 – 1.76 (m, 3H), 1.60 (d, $J$ = 9.4 Hz, 3H), 1.55 – 1.35 (m, 9H), 1.30 (tt, $J$ = 12.8, 5.6 Hz, 9H), 1.24 – 1.11 (m, 3H), 1.10 (s, 4H), 1.04 – 0.98 (m, 1H), 0.96 (s, 3H), 0.94 (d, $J$ = 6.5 Hz, 3H), 0.69 (s, 3H). $^{13}$C NMR (151 MHz, Methanol-$d_4$) δ 178.0, 170.7, 135.4, 130.3, 129.9, 128.7, 128.1, 59.1, 58.6, 57.7, 57.4, 43.2, 41.6, 41.3, 37.0, 36.9, 36.5, 35.8, 35.5, 32.2, 31.8, 30.6, 29.0, 27.8, 27.3, 25.0, 24.4, 23.4, 21.7, 18.6, 12.3.

*Synthesis of N-acetylcadaverine-3-oxo-lithocholic acid (N-acetylcadaverine-3-lithocholic acid):* To a solution of 3-oxolithocholic acid (300 mg, 1 equiv, 801 μmol) in methanol (20 mL) was added acetic acid (458 μL, 10 equiv, 8.01 mmol) followed by N-acetylcadaverine (347 mg, 3 equiv, 2.40 mmol). The mixture was stirred at room temperature for 3 h. Sodium cyanoborohydride (151 mg, 3 equiv, 2.40 mmol) was then added, and the reaction was continued for a further 16 h at room temperature. Reaction progress was monitored by LC-MS; after complete consumption of the starting material the solvent was removed under reduced pressure. The resulting crude material was redissolved in methanol and purified by reverse-phase C18 chromatography, employing a linear gradient of 50 %–70 % aqueous acetonitrile (0.1 % TFA). Two diastereomeric fractions were collected and concentrated, affording:

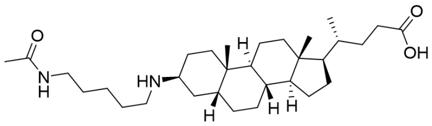

N-acetylcadaverine-3(S)-lithocholic acid, 75 mg, 19 % yield. $^{1}$H NMR (600 MHz, Methanol-$d_4$) δ 3.46 (s, 1H), 3.18 (t, $J$ = 7.0 Hz, 2H), 3.01 – 2.96 (m, 2H), 2.40 – 2.16 (m, 4H), 2.03 – 1.99 (m, 1H), 1.99 – 1.94 (m, 1H), 1.93 (s, 2H), 1.90 (dd, $J$ = 8.5, 4.5 Hz, 1H), 1.72 (ddd, $J$ = 15.8, 11.9, 5.6 Hz, 5H), 1.57 (ddt, $J$ = 32.8, 11.1, 6.6 Hz, 6H), 1.51 – 1.39 (m, 9H), 1.39 – 1.25 (m, 6H), 1.25 – 1.20 (m, 2H), 1.14 (dtt, $J$ = 21.3, 11.2, 5.1 Hz, 4H), 1.04 (s, 2H), 0.94 (t, $J$ = 7.1 Hz, 2H), 0.70 (d, $J$ = 3.5 Hz, 3H). $^{13}$C NMR (151 MHz, Methanol-$d_4$) δ 178.2, 173.4, 49.6, 45.5, 43.9, 41.3, 39.9, 36.0, 32.3, 32.2, 32.0, 31.8, 30.7, 29.9, 29.2, 28.5, 27.4, 27.0, 26.5, 25.2, 24.8, 22.6, 22.0.

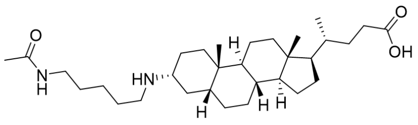

N-acetylcadaverine-3(R)-lithocholic acid, 70 mg, 17 % yield. $^{1}$H NMR (600 MHz, Methanol-

*$d_4$*) δ 3.18 (t, *J* = 7.1 Hz, 2H), 3.10 (ddt, *J* = 12.0, 8.7, 4.2 Hz, 2H), 3.03 – 2.97 (m, 2H), 2.34 (dtd, *J* = 20.6, 9.9, 4.8 Hz, 1H), 2.22 (tdd, *J* = 15.6, 9.3, 4.7 Hz, 2H), 2.08 – 1.95 (m, 4H), 1.93 (s, 3H), 1.89 (dd, *J* = 15.3, 9.4 Hz, 3H), 1.80 (dtd, *J* = 16.7, 6.5, 3.0 Hz, 4H), 1.68 (pent, *J* = 7.8 Hz, 5H), 1.64 – 1.58 (m, 1H), 1.55 (pent, *J* = 6.5 Hz, 2H), 1.53 – 1.46 (m, 6H), 1.38 – 1.26 (m, 7H), 1.15 – 1.05 (m, 9H), 1.00 (s, 3H), 0.94 (t, *J* = 7.1 Hz, 3H), 0.70 (d, *J* = 3.4 Hz, 3H). $^{13}$C NMR (151 MHz, Methanol-*$d_4$*) δ 178.1, 173.4, 59.1, 57.9, 57.6, 49.6, 45.7, 43.9, 43.2, 41.8, 41.5, 39.9, 37.1, 36.7, 35.9, 35.8, 32.3, 32.0, 30.8, 29.9, 29.2, 27.9, 27.4, 27.1, 25.2, 25.0, 24.8, 23.7, 22.6, 21.9, 18.7, 12.4.

*Synthesis of Phenylalanine_3-oxo-deoxycholic acid (N-[3-deoxycholyl]phenylalanine)*:
To a stirred solution 3-oxo-deoxycholic acid of (100 mg, 256 μmol, 1.0 equiv) in methanol (5.0 mL) were added acetic acid (147 μL, 2.56 mmol, 10 equiv) and L-phenylalanine (127 mg, 768 μmol, 3.0 equiv). The reaction mixture was stirred at room temperature for 2 h. Sodium cyanoborohydride (48.3 mg, 768 μmol, 3.0 equiv) was then added, and the mixture was stirred at room temperature overnight. The reaction progress was monitored by LC–MS. Upon completion, the solvent was removed under reduced pressure to remove methanol and excess acetic acid. The crude residue was diluted with a mixture of acetonitrile/water and purified by reverse-phase flash chromatography on a C18 column, eluting with 50–60% acetonitrile in water containing 0.1% TFA as a gradient. Fractions containing the desired products were combined and concentrated to afford the following diastereomers:

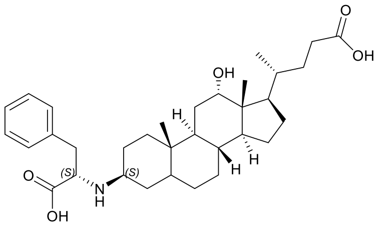


*N*-[3(S)-deoxycholyl]phenylalanine (27 mg, 20%). $^1$H NMR (600 MHz, Methanol-*$d_4$*) δ 7.47 – 7.29 (m, 5H), 4.27 (t, *J* = 7.6 Hz, 1H), 4.00 (t, *J* = 2.7 Hz, 1H), 3.58 (s, 1H), 3.41 – 3.35 (m, 2H), 3.29 (dd, *J* = 13.8, 8.5 Hz, 1H), 2.37 (ddd, *J* = 15.2, 9.8, 5.2 Hz, 1H), 2.30 – 2.19 (m, 3H), 1.92 (dtt, *J* = 18.8, 9.5, 4.0 Hz, 4H), 1.88 – 1.79 (m, 4H), 1.79 – 1.72 (m, 2H), 1.64 (ddd, *J* = 23.1, 17.4, 12.2 Hz, 6H), 1.58 – 1.52 (m, 2H), 1.51 – 1.42 (m, 6H), 1.35 (ddt, *J* = 20.4, 6.7, 4.4 Hz, 5H), 1.27 – 1.17 (m, 3H), 1.12 (qd, *J* = 16.0, 7.5 Hz, 3H), 1.03 (d, *J* = 6.6 Hz, 2H), 1.01 (s, 2H), 0.74 (s, 2H). $^{13}$C NMR -DEPT-135 (151 MHz, Methanol-*$d_4$*) δ 130.2, 129.9, 128.7, 73.6, 73.3, 72.2, 67.9, 61.9, 61.2, 57.0, 49.0, 47.9, 43.5, 37.3, 36.5, 34.2, 32.0, 31.7, 30.6, 29.6, 28.4, 27.8, 27.1, 26.6, 24.5, 23.2, 23.0, 17.3, 12.9.

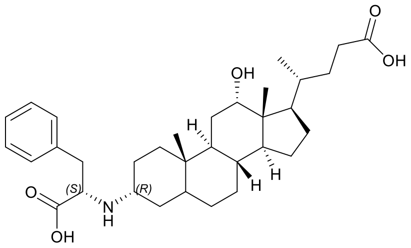


*N*-[3(R)-deoxycholyl]phenylalanine (26 mg, 19%). $^1$H NMR (600 MHz, Methanol-*$d_4$*) δ 7.33 (tt, *J* = 15.3, 7.1 Hz, 5H), 4.35 (t, *J* = 6.7 Hz, 1H), 3.29 – 3.20 (m, 2H), 3.13 (t, *J* = 11.9 Hz,

1H), 2.34 (ddd, *J* = 15.1, 9.8, 5.2 Hz, 1H), 2.22 (ddd, *J* = 15.7, 9.4, 6.8 Hz, 1H), 2.00 – 1.72 (m, 11H), 1.69 – 1.25 (m, 22H), 1.18 – 1.04 (m, 3H), 1.01 (d, *J* = 6.5 Hz, 3H), 0.99 (dd, *J* = 5.4, 2.6 Hz, 1H), 0.97 – 0.95 (m, 1H), 0.71 (s, 2H). $^{13}$C NMR (151 MHz, Methanol-$d_4$) δ 129.0, 128.6, 127.4, 72.4, 72.2, 71.1, 66.8, 60.8, 58.0, 57.5, 46.7, 42.3, 42.0, 35.9, 35.7, 35.4, 34.5, 33.5, 30.9, 30.6, 29.4, 28.4, 27.3, 26.6, 25.9, 23.4, 23.3, 22.0, 16.2, 11.8.

*Synthesis of 5-ASA_pyruvate (5-((1-carboxyethyl)amino)-2-hydroxybenzoic acid)*:
To a stirred solution of 2-oxopropanoic acid (500 mg, 5.68 mmol, 1.0 equiv) in methanol (20 mL) were added acetic acid (3.25 mL, 56.8 mmol, 10 equiv) and 5-amino-2-hydroxybenzoic acid (2.61 g, 17.0 mmol, 3.0 equiv). The reaction mixture was stirred at room temperature for 2 h. Sodium cyanoborohydride (1.07 g, 17.0 mmol, 3.0 equiv) was then added, and the mixture was stirred at room temperature overnight. Reaction progress was monitored by LC–MS. Upon completion, the solvent was removed under reduced pressure to remove methanol and excess acetic acid. The crude residue was diluted with a mixture of acetonitrile/water and purified by reverse-phase flash chromatography on a C18 column, eluting with 0–30% acetonitrile/water as a gradient. Fractions containing the desired product were combined and concentrated to afford 5-((1-carboxyethyl)amino)-2-hydroxybenzoic acid (145 mg, 11.3% yield);

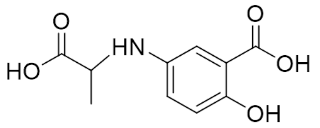


$^1$H NMR (600 MHz, Methanol-$d_4$) δ 7.20 (d, *J* = 2.9 Hz, 1H), 6.97 (dd, *J* = 8.8, 2.9 Hz, 1H), 6.78 (d, *J* = 8.8 Hz, 1H), 3.98 (q, *J* = 7.0 Hz, 1H), 1.46 (d, *J* = 7.0 Hz, 3H). $^{13}$C NMR (151 MHz, Methanol-$d_4$) δ 177.98, 173.45, 156.72, 139.88, 139.84, 124.90, 118.68, 115.76, 55.00, 18.49.

*Synthesis of 5-ASA_ribose (2-hydroxy-5-(((2S,3S,4R)-2,3,4,5-tetrahydroxypentyl)amino)benzoic acid):* To a stirred solution of D-Ribose (300 mg, 1 Eq, 2.00 mmol) in MeOH (16 g, 20 mL) were added acetic acid (1.20 g, 1.14 mL, 10 Eq, 20.0 mmol) and 5-amino-2-hydroxybenzoic acid (918 mg, 3 Eq, 5.99 mmol). The reaction mixture was stirred at room temperature for 2 h. Sodium cyanoborohydride (377 mg, 3 Eq, 5.99 mmol) was then added, and the mixture was stirred at room temperature overnight. The reaction progress was monitored by LC–MS. Upon completion, the solvent was removed under reduced pressure to remove methanol and excess acetic acid. The crude residue was diluted with a mixture of acetonitrile/water and purified by reverse-phase flash chromatography on a C18 column, eluting with 50–60% acetonitrile in water containing 0.1% TFA as a gradient. Fractions containing the desired products were combined and concentrated to afford the 2-hydroxy-5-(((2S,3S,4R)-2,3,4,5-tetrahydroxypentyl)amino)benzoic acid (220 mg, 766 µmol, 38.3 %).

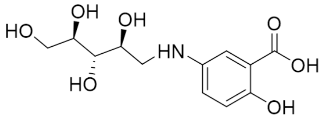


$^1$H NMR (600 MHz, Methanol-$d_4$) δ 8.01 (d, *J* = 6.3 Hz, 1H), 7.61 (dt, *J* = 6.4, 3.3 Hz, 1H), 7.10 (t, *J* = 11.6 Hz, 1H), 4.05 (dd, *J* = 8.1, 3.9 Hz, 1H), 3.77 – 3.71 (m, 1H), 3.71 – 3.65 (m, 2H), 3.64 – 3.59 (m, 1H), 3.58 – 3.45 (m, 2H). $^{13}$C NMR (151 MHz, Methanol-$d_4$) δ 171.0,

161.6, 129.2, 128.6, 127.5, 124.2, 123.7, 118.7, 114.1, 73.2, 72.6, 67.1, 63.2, 53.5.

*Synthesis of sulfapyridine_glyceraldehyde (4-((2,3-dihydroxypropyl)amino)-N-(pyridin-2-yl)benzenesulfonamide)*: To a stirred solution of 2,3-Dihydroxypropanal (200 mg, 1 Eq, 2.22 mmol) in methanol (20 mL) were added Sulfapyridine (1.66 g, 3 Eq, 6.66 mmol) and acetic acid (1.33 g, 10 Eq, 22.2 mmol). The reaction mixture was stirred at room temperature for 2 h. Sodium cyanoborohydride (419 mg, 3 Eq, 6.66 mmol) was then added, and the mixture was stirred at room temperature overnight. The reaction progress was monitored by LC–MS. Upon completion, the solvent was removed under reduced pressure to remove methanol and excess acetic acid. The crude residue was diluted with a mixture of acetonitrile/water and purified by reverse-phase flash chromatography on a C18 column, eluting with 50–60% acetonitrile in water containing 0.1% TFA as a gradient. Fractions containing the desired products were combined and concentrated to afford the 4-((2,3-dihydroxypropyl)amino)-N-(pyridin-2-yl)benzenesulfonamide (210 mg, 29.2 %).

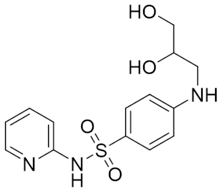


$^1$H NMR (600 MHz, Methanol-$d_4$) δ 8.18 (d, $J$ = 5.1 Hz, 1H), 7.96 – 7.88 (m, 1H), 7.69 (d, $J$ = 8.9 Hz, 1H), 7.32 (d, $J$ = 8.5 Hz, 1H), 7.17 – 7.11 (m, 1H), 6.71 (d, $J$ = 8.9 Hz, 1H), 3.82 (dq, $J$ = 10.1, 5.2 Hz, 1H), 3.63 – 3.54 (m, 2H), 3.36 (d, $J$ = 4.6 Hz, 1H), 3.16 (dd, $J$ = 13.4, 7.1 Hz, 1H). $^{13}$C NMR (151 MHz, Methanol-$d_4$) δ 154.37, 152.17, 143.94, 143.66, 130.26, 126.40, 119.09, 115.77, 112.76, 71.51, 65.27, 46.94.

*Synthesis of sulfapyridine_2-oxoglutaric acid ((4-(N-(pyridin-2-yl)sulfamoyl)phenyl)glutamic acid)*: To a stirred solution of alpha-Ketoglutaric acid (200 mg, 1 Eq, 1.37 mmol) in methanol (10 mL) were added Sulfapyridine (1.02 g, 3 Eq, 4.11 mmol) and acetic acid (822 mg, 10 Eq, 13.7 mmol). The reaction mixture was stirred at room temperature for 2 h. Sodium cyanoborohydride (258 mg, 3 Eq, 4.11 mmol) was then added, and the mixture was stirred at room temperature overnight. The reaction progress was monitored by LC–MS. Upon completion, the solvent was removed under reduced pressure to remove methanol and excess acetic acid. The crude residue was diluted with a mixture of acetonitrile/water and purified by reverse-phase flash chromatography on a C18 column, eluting with 50–60% acetonitrile in water containing 0.1% TFA as a gradient. Fractions containing the desired products were combined and concentrated to afford the (4-(N-(pyridin-2-yl)sulfamoyl)phenyl)glutamic acid (210 mg, 554 μmol, 40.4 %);

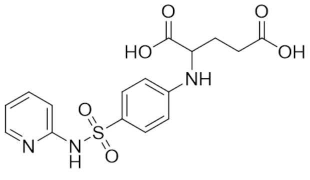

$^{1}$H NMR (600 MHz, DMSO-$d_6$) δ 11.0 -13.0 (bs, 2H), 8.09 (d, $J$ = 4.3 Hz, 1H), 7.76 – 7.63 (m, 1H), 7.60 (d, $J$ = 9.0 Hz, 2H), 7.09 (d, $J$ = 8.5 Hz, 1H), 6.93 – 6.85 (m, 1H), 6.75 (s, 1H), 6.61 (d, $J$ = 8.9 Hz, 2H), 4.00 (dt, $J$ = 11.2, 5.6 Hz, 1H), 2.39 – 2.35 (m, 2H), 2.03 (dq, $J$ = 13.7, 7.5 Hz, 1H), 1.90 (dq, $J$ = 14.9, 8.4 Hz, 1H). $^{13}$C NMR (151 MHz, DMSO-$d_6$) δ 174.1, 173.8, 152.4, 151.3, 141.8, 138.9, 128.7, 127.5, 119.5, 112.3, 111.2, 54.4, 30.1, 27.0.

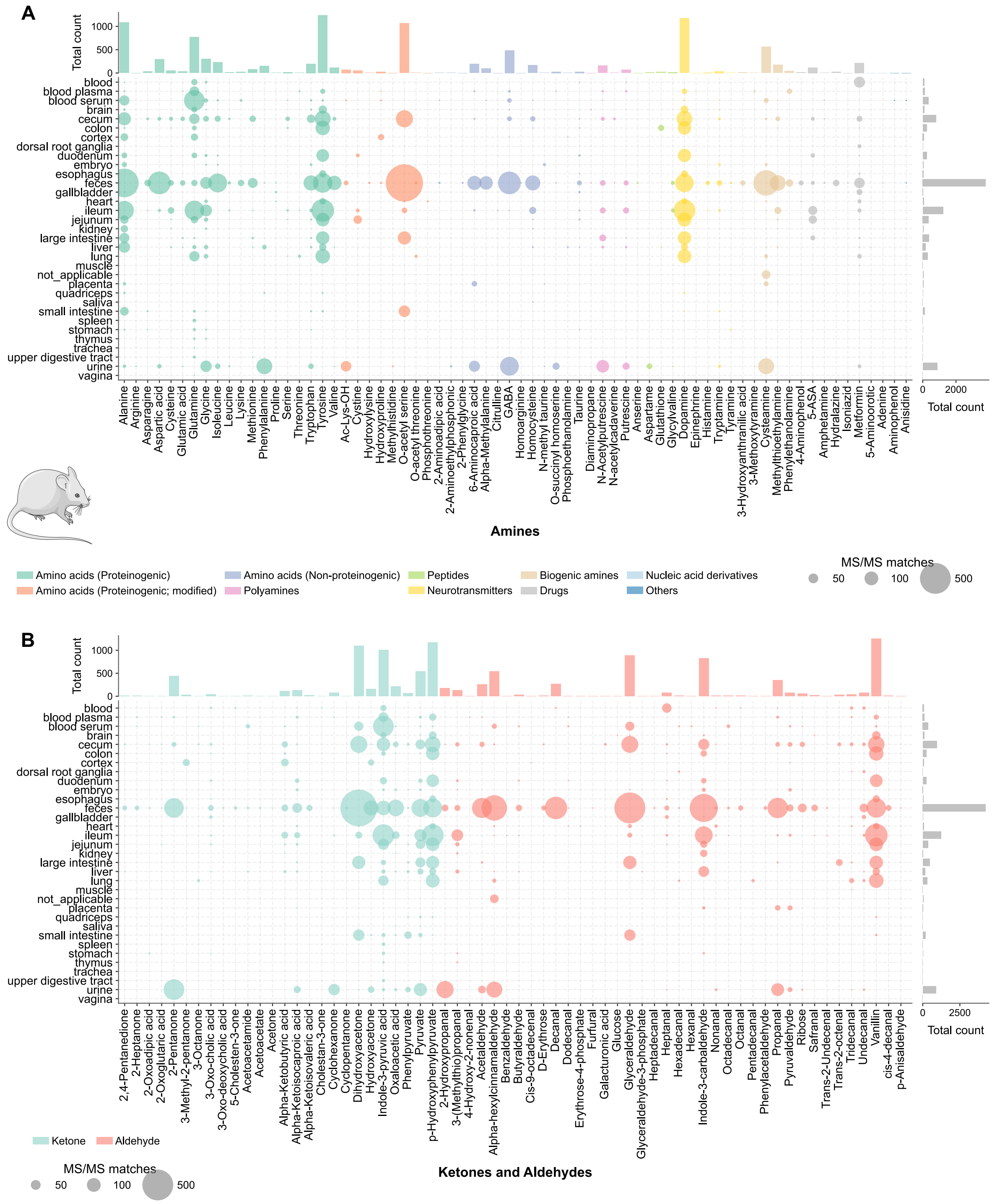


**Fig. S1. Pan-repository MS/MS matches and their organ and biofluid distributions in rodents. (A)** Distribution of MS/MS of alkamines synthesized from amines in rodent samples. **(B)** Organ and biofluid distribution of MS/MS of alkamines synthesized from aldehydes and ketones in rodent samples.

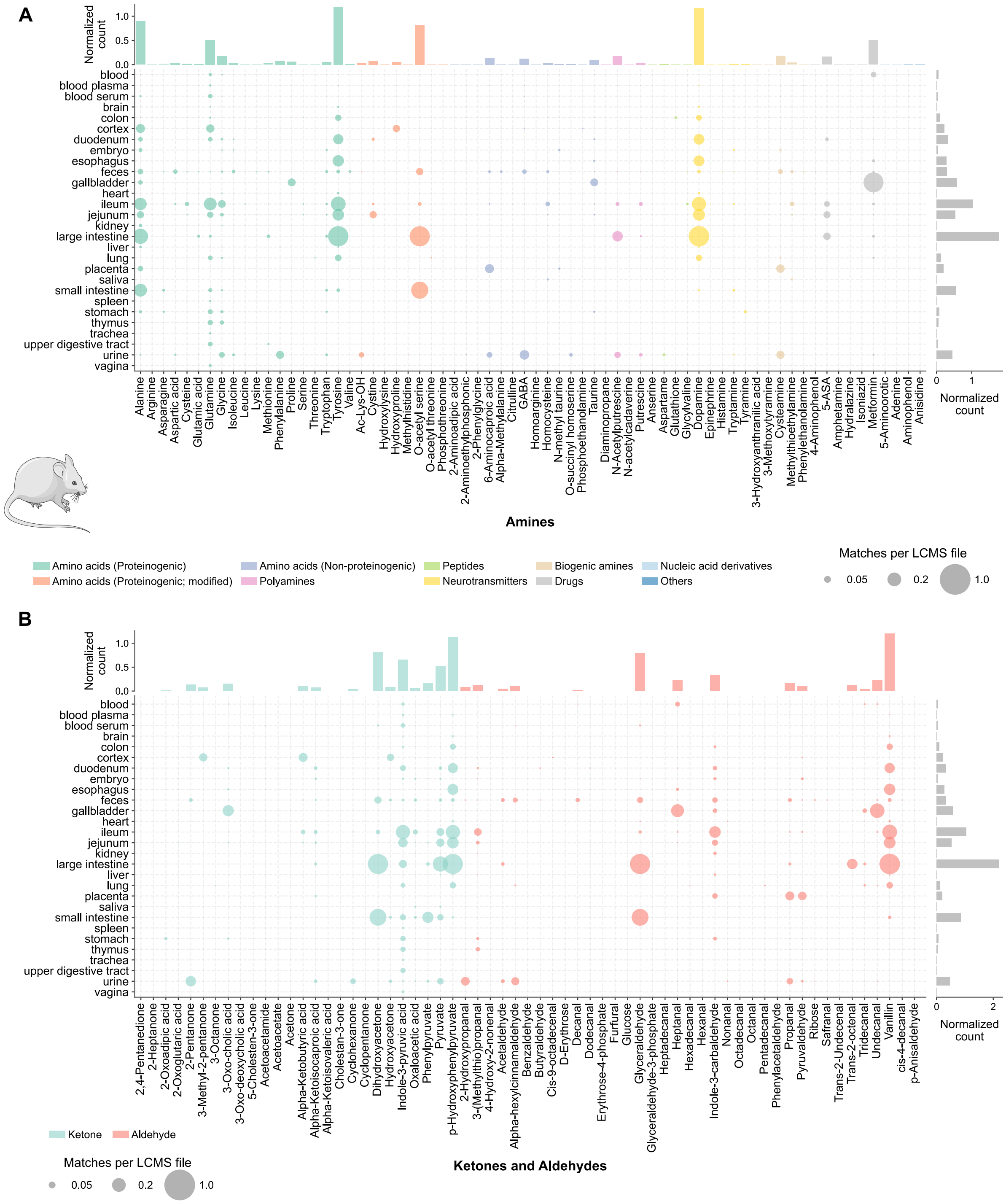


**Fig. S2. Normalizes by the number of sample types of Pan-repository MS/MS matches and their organ and biofluid distributions in rodent samples. (A)** Distribution of MS/MS of alkamines synthesized from amines in rodent samples normalized by the number of sample types. **(B)** Organ and biofluid distribution of MS/MS of alkamines synthesized from aldehydes and ketones in rodent samples normalized by the number of sample types.

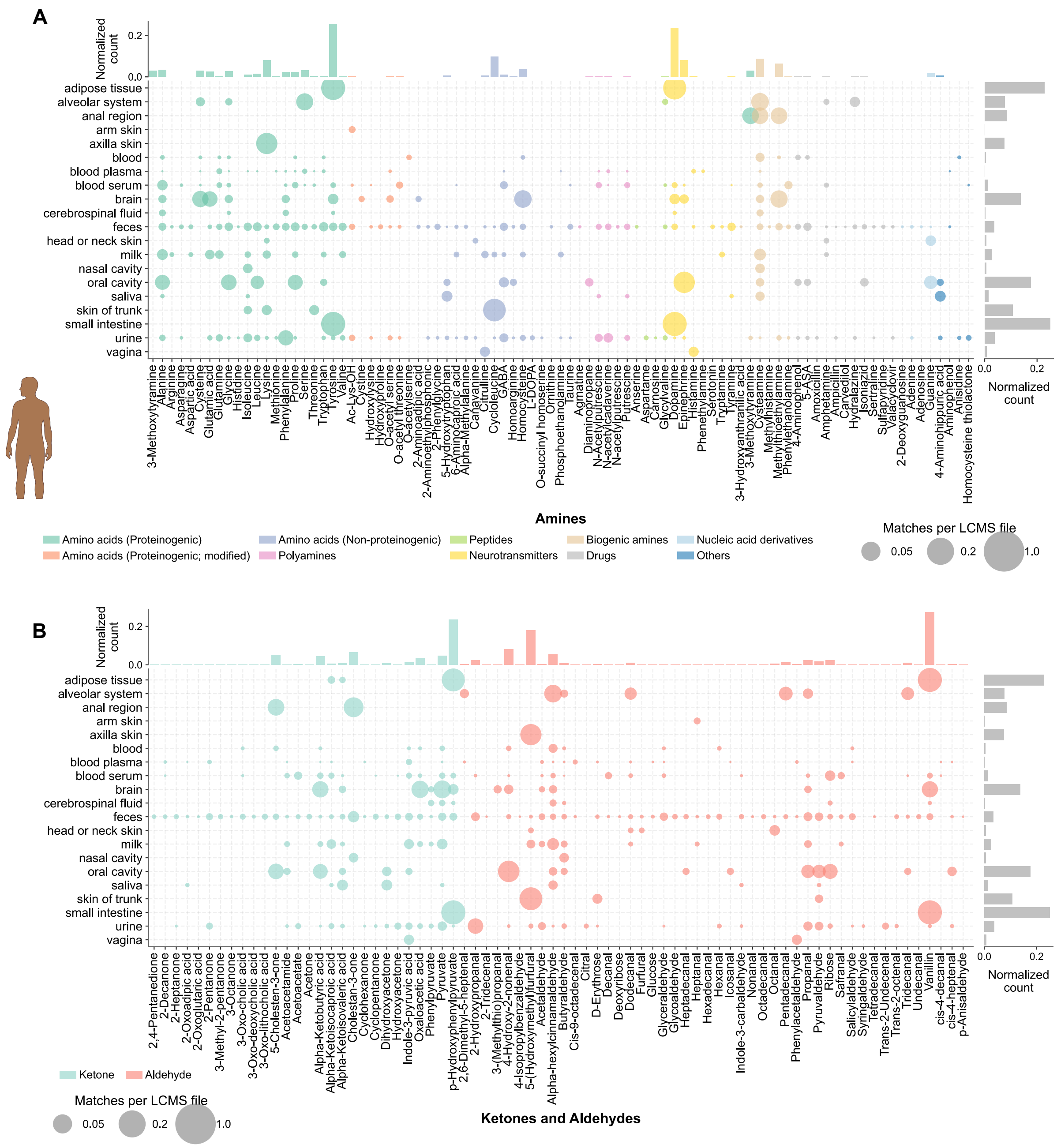


**Fig. S3. Normalizes by the number of sample types of Pan-repository MS/MS matches and their organ and biofluid distributions in human samples. (A)** Distribution of MS/MS of alkamines synthesized from amines in human samples normalized by the number of sample types. **(B)** Organ and biofluid distribution of MS/MS of alkamines synthesized from aldehydes and ketones in human samples normalized by the number of sample types.

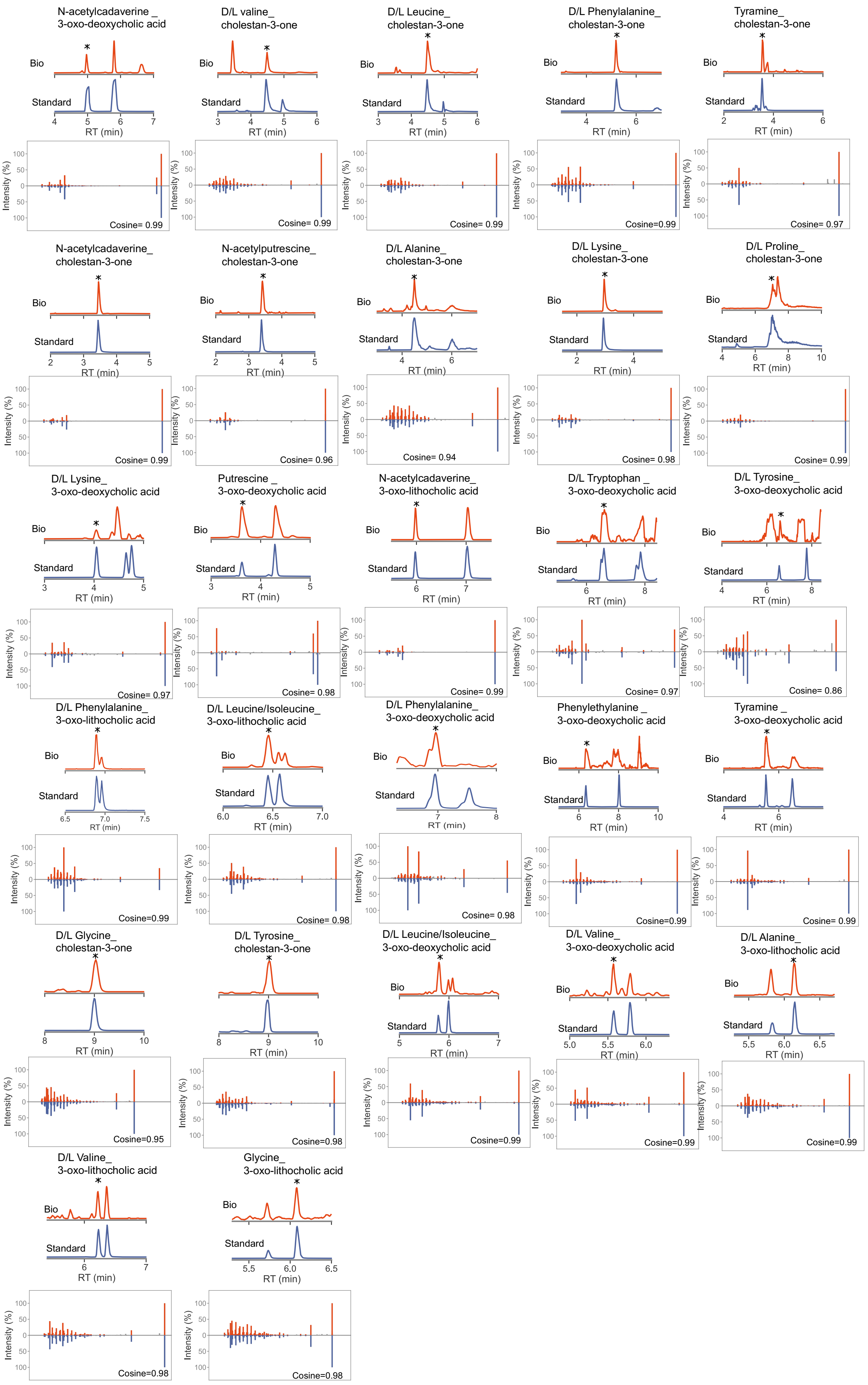
N-acetylcadaverine _ 3-oxo-deoxycholic acid
Bio
Standard
RT (min)
Intensity (%)
Cosine= 0.99
D/L valine_ cholestan-3-one
Cosine= 0.99
D/L Leucine_ cholestan-3-one
Cosine= 0.99
D/L Phenylalanine_ cholestan-3-one
Cosine=0.99
Tyramine_ cholestan-3-one
Cosine= 0.97
N-acetylcadaverine_ cholestan-3-one
Cosine= 0.99
N-acetylputrescine_ cholestan-3-one
Cosine= 0.96
D/L Alanine_ cholestan-3-one
Cosine= 0.94
D/L Lysine_ cholestan-3-one
Cosine= 0.98
D/L Proline_ cholestan-3-one
Cosine= 0.99
D/L Lysine_ 3-oxo-deoxycholic acid
Cosine= 0.97
Putrescine _ 3-oxo-deoxycholic acid
Cosine= 0.98
N-acetylcadaverine_ 3-oxo-lithocholic acid
Cosine= 0.99
D/L Tryptophan _ 3-oxo-deoxycholic acid
Cosine= 0.97
D/L Tyrosine_ 3-oxo-deoxycholic acid
Cosine= 0.86
D/L Phenylalanine_ 3-oxo-lithocholic acid
Cosine=0.99
D/L Leucine/Isoleucine_ 3-oxo-lithocholic acid
Cosine= 0.98
D/L Phenylalanine_ 3-oxo-deoxycholic acid
Cosine= 0.98
Phenylethylamine _ 3-oxo-deoxycholic acid
Cosine=0.99
Tyramine _ 3-oxo-deoxycholic acid
Cosine= 0.99
D/L Glycine_ cholestan-3-one
Cosine=0.95
D/L Tyrosine_ cholestan-3-one
Cosine=0.98
D/L Leucine/Isoleucine_ 3-oxo-deoxycholic acid
Cosine=0.99
D/L Valine_ 3-oxo-deoxycholic acid
Cosine=0.99
D/L Alanine_ 3-oxo-lithocholic acid
Cosine=0.99
D/L Valine_ 3-oxo-lithocholic acid
Cosine=0.98
Glycine_ 3-oxo-lithocholic acid
Cosine=0.98

**Fig. S4. Structure validation of synthesized bile acid and steroid alkamines with biological samples using chromatography and MS/MS.** Chromatographic comigration of bile acids and steroids alkamines including 3-oxo-lithocholic acid, 3-oxo-deoxycholic and cholestan-3-one synthesized alkamines conjugates with fecal samples from human cohorts and feline (African lion and cheetah) using LC-MS/MS condition. The asterisk represents the MS/MS of the matched peak from the biological with the synthetic standard that is shown.

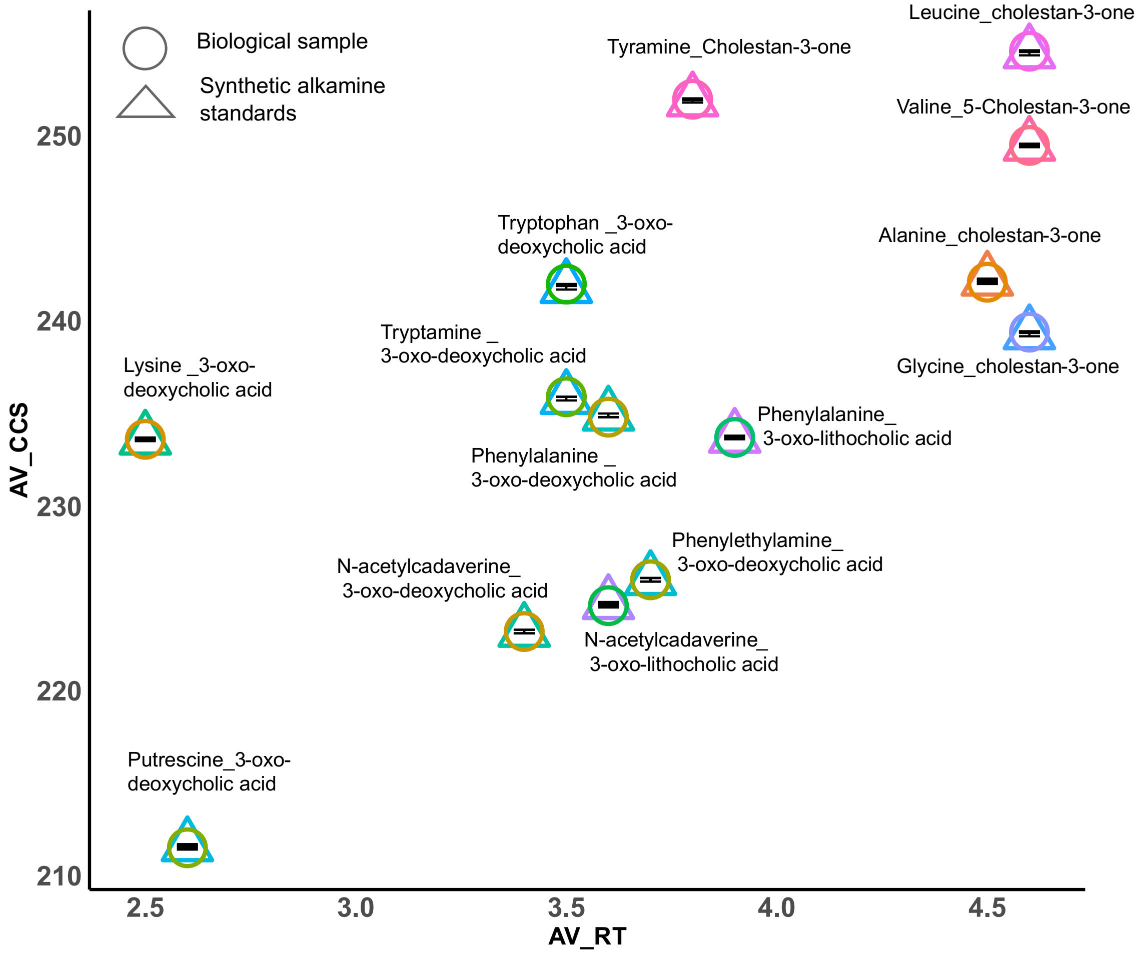


**Fig. S5. RT and ion mobility matching synthesized bile acid and steroid alkamines with biological samples.** Collision cross section comigration of bile acids and steroids alkamines including 3-oxo-lithocholic acid, 3-oxo-deoxycholic and cholestan-3-one synthesized alkamines conjugates with fecal samples from human cohorts and feline (African lion and cheetah) using ion mobility.

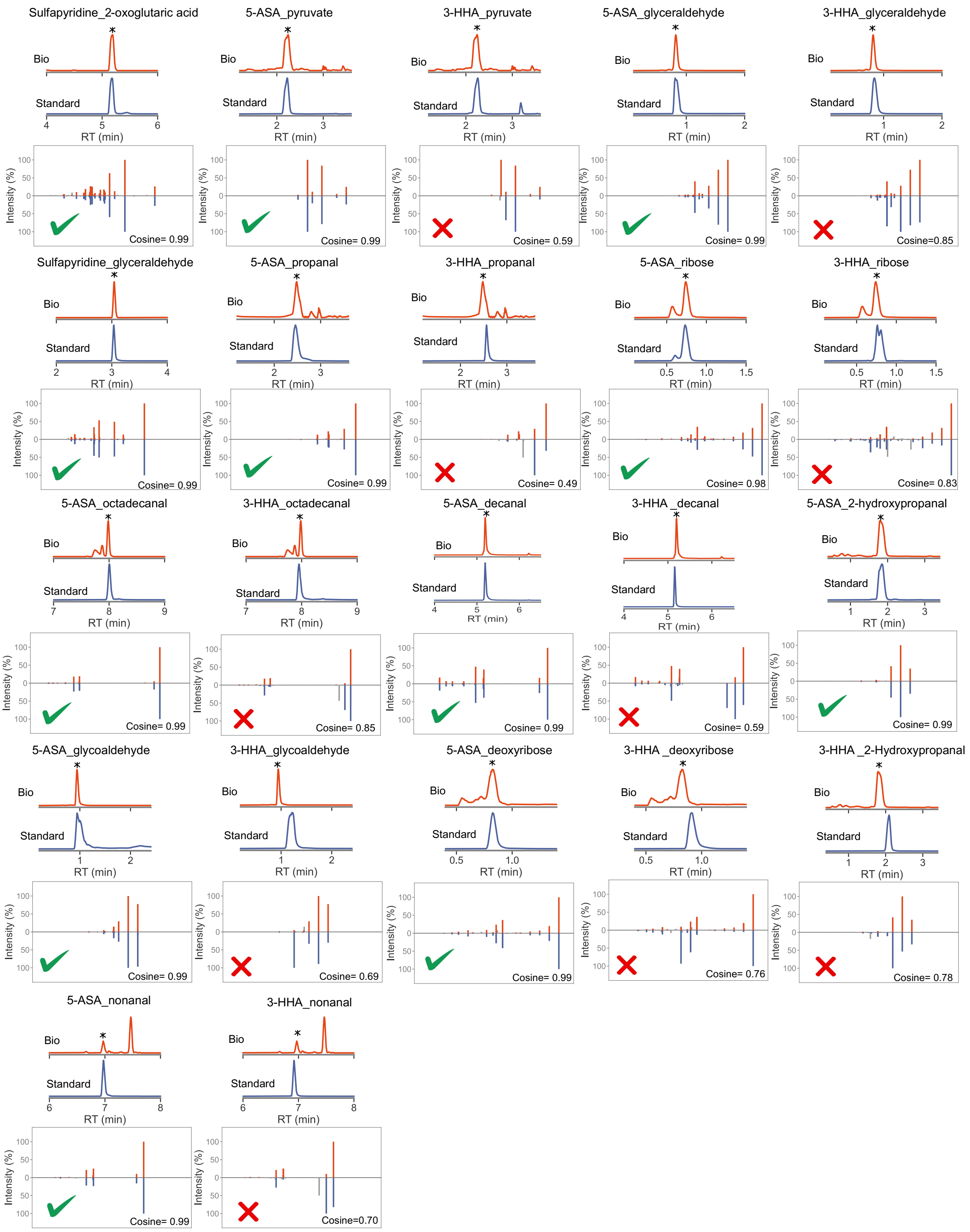


**Fig. S6. Structure validation of synthesized drug conjugate alkamines with biological samples using chromatography and MS/MS.** Chromatographic comigration of 5-aminosalicylic acid (5-ASA) 3-hydroxyanthranillic acid (3-HHA) and sulfapyridine synthesized alkamines conjugates with human fecal samples of Rheumatoids arthritis cohort. The asterisk represents the matched peak from the biological with the synthetic standard. And the correct symbol representing the correct matched peak.

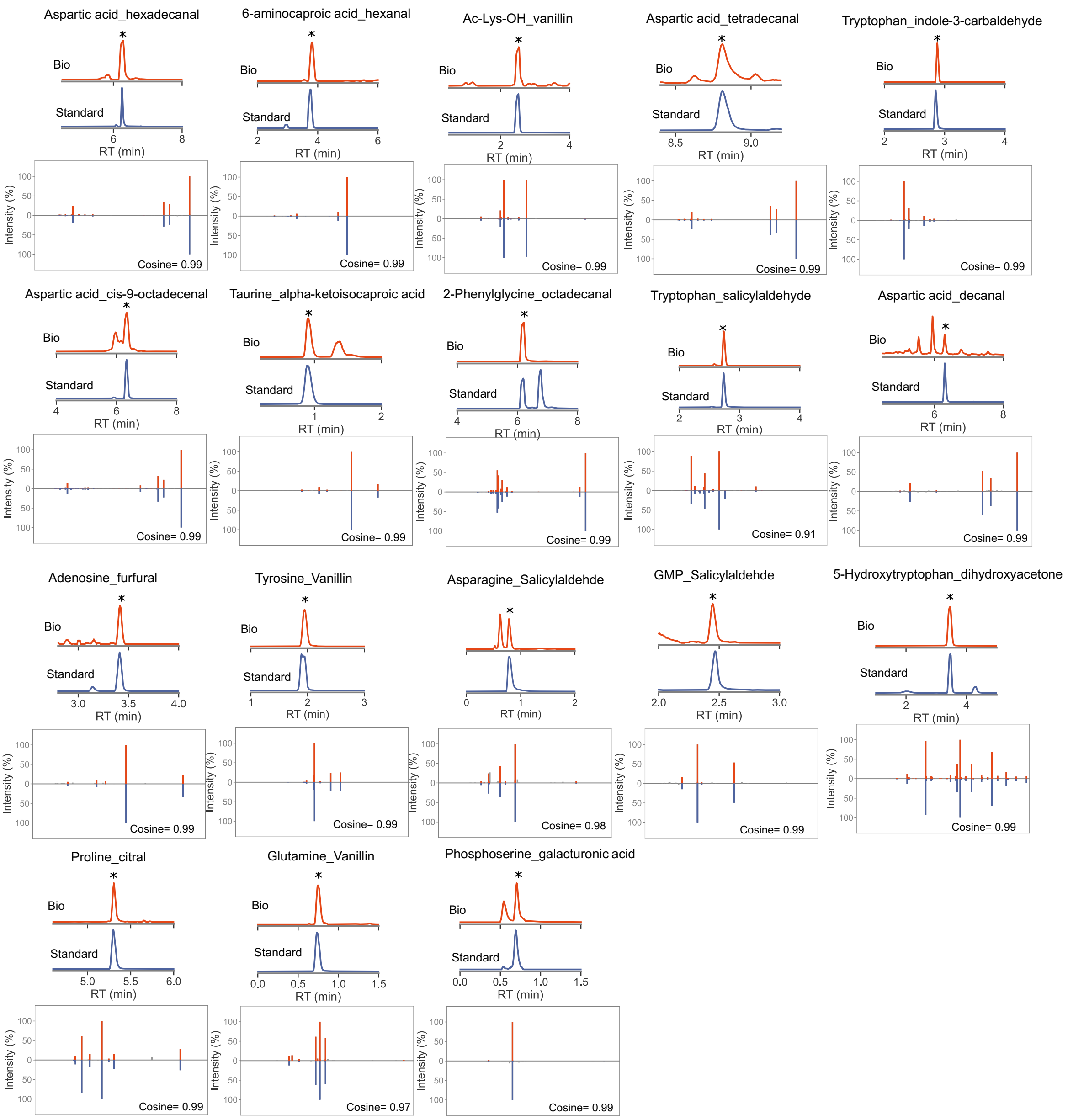


**Fig. S7. Structure validation of synthesized alkamines with biological samples using chromatography and MS/MS.** Chromatographic comigration of newly synthesized alkamines with fecal samples from lizards and food extracts. The asterisk represents the MS/MS matched peak from the biological with the synthetic standard shown.

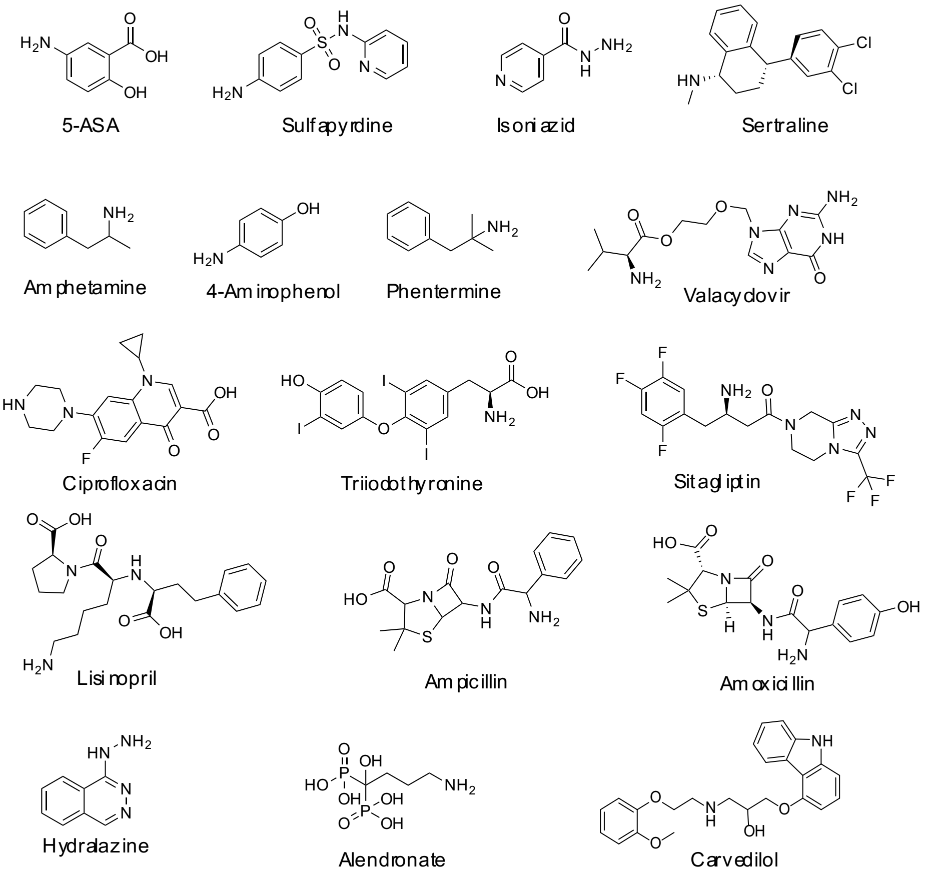


**Fig. S8. Drug alkamines were generated from seventeen amine-containing drugs through reactions with aldehydes and ketones.** Structures of 17 amine containing drugs reacted with aldehydes and ketones to produce drug alkamines.

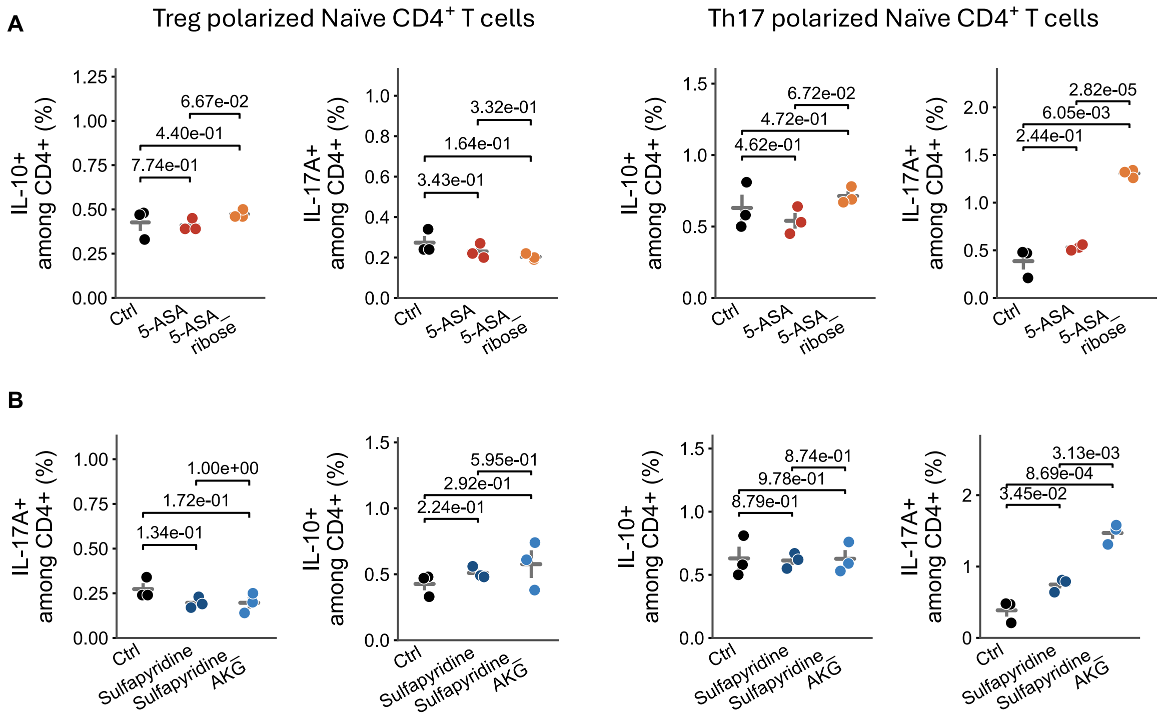


**Fig. S9. Immune Naïve CD4+ T cells activity when treated with 5-ASA_ribose and Sulfapyridine_Alphaketoglutarate**. **(A)** Naïve CD4+ T cells were polarized under Th17 or Treg conditions and treated with 50 µM 5-ASA and 5-ASA_ribose, and compared to vehicle control (DMSO, Contr). **(B)** Naïve CD4+ T cells polarized under Th17 or Treg conditions were treated with 50 µM sulfapyridine and sulfapyridine_alpha ketoglutarate, and compared to vehicle control (DMSO, Contr).

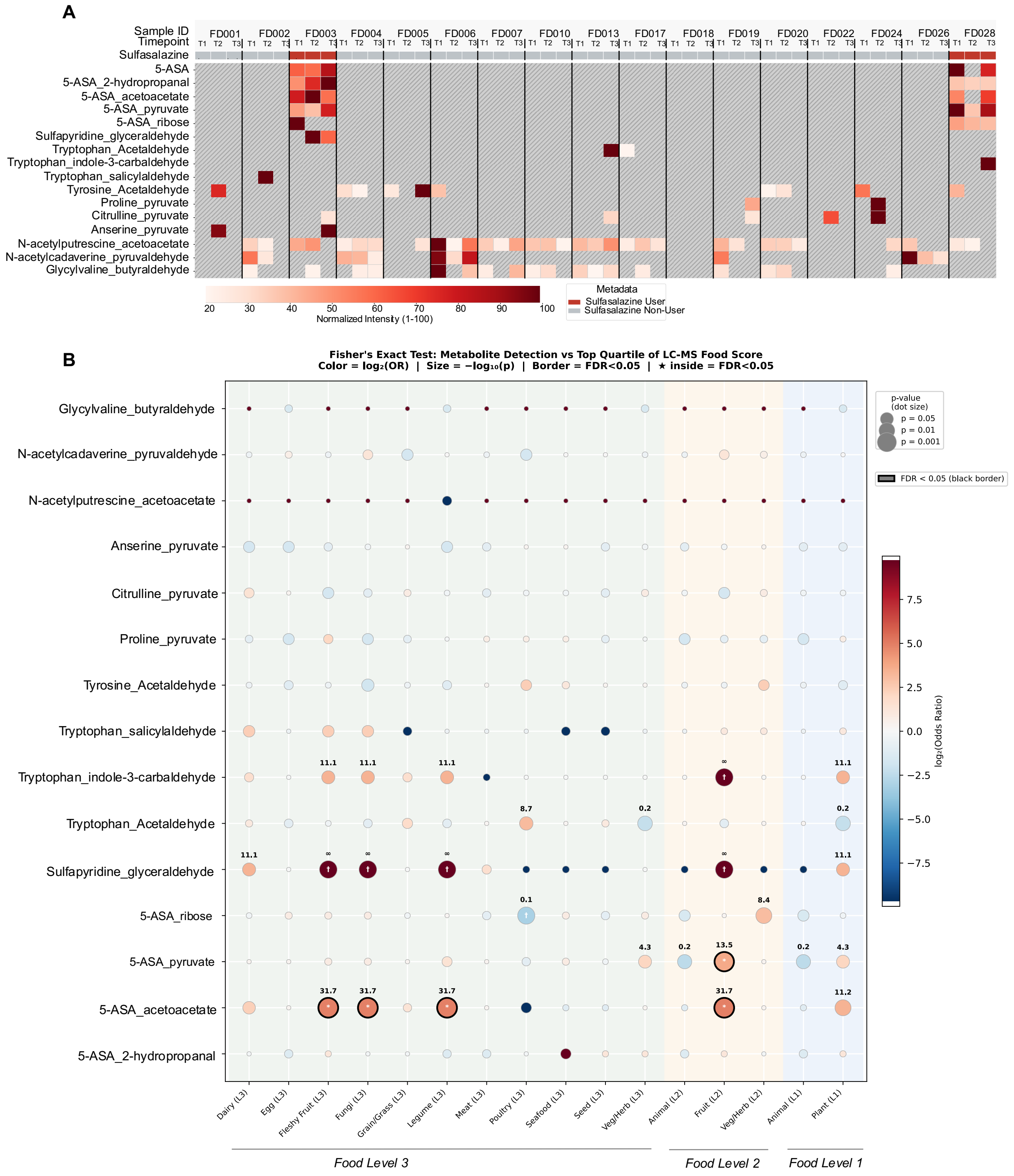


**Fig. S10. Overview of levels of alkamines, including drug conjugates detected in Rheumatoid arthritis cohort in relation to diet intake**. **(A)** Heatmap showing levels of alkamines across Rheumatoid arthritis cohort. **(B)** Fisher exact test was performed to compare detection of alkylamine conjugate with the individuals in the top quartile of food scores obtained from metabolomics data(*12*). Each bubble represents one test; dot size encodes statistical significance as $-\log_{10}$($p$-value) and dot color encodes effect size as $\log_2$(odds ratio), ranging from deep red (strong positive association) to deep blue (strong negative association). Dots with a black border indicate FDR-corrected $q < 0.05$ (Benjamini–Hochberg).

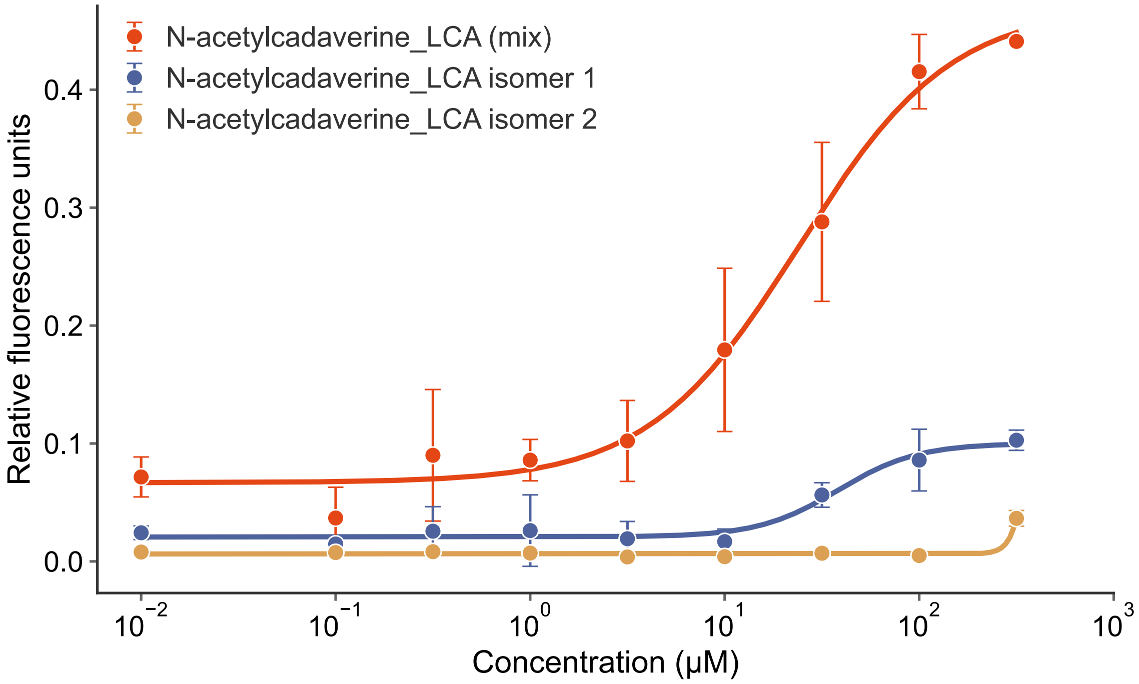


**Fig. S11. Dose response curve for *N*-acetylcadaverine_LCA mix, isomer1 (S) and isomer2 (R).** The dose response curve for *N*-acetylcadaverine_LCA mix isomers and in comparison, with the isomer1 (S) and isomer2, data availability **(Table S14).**

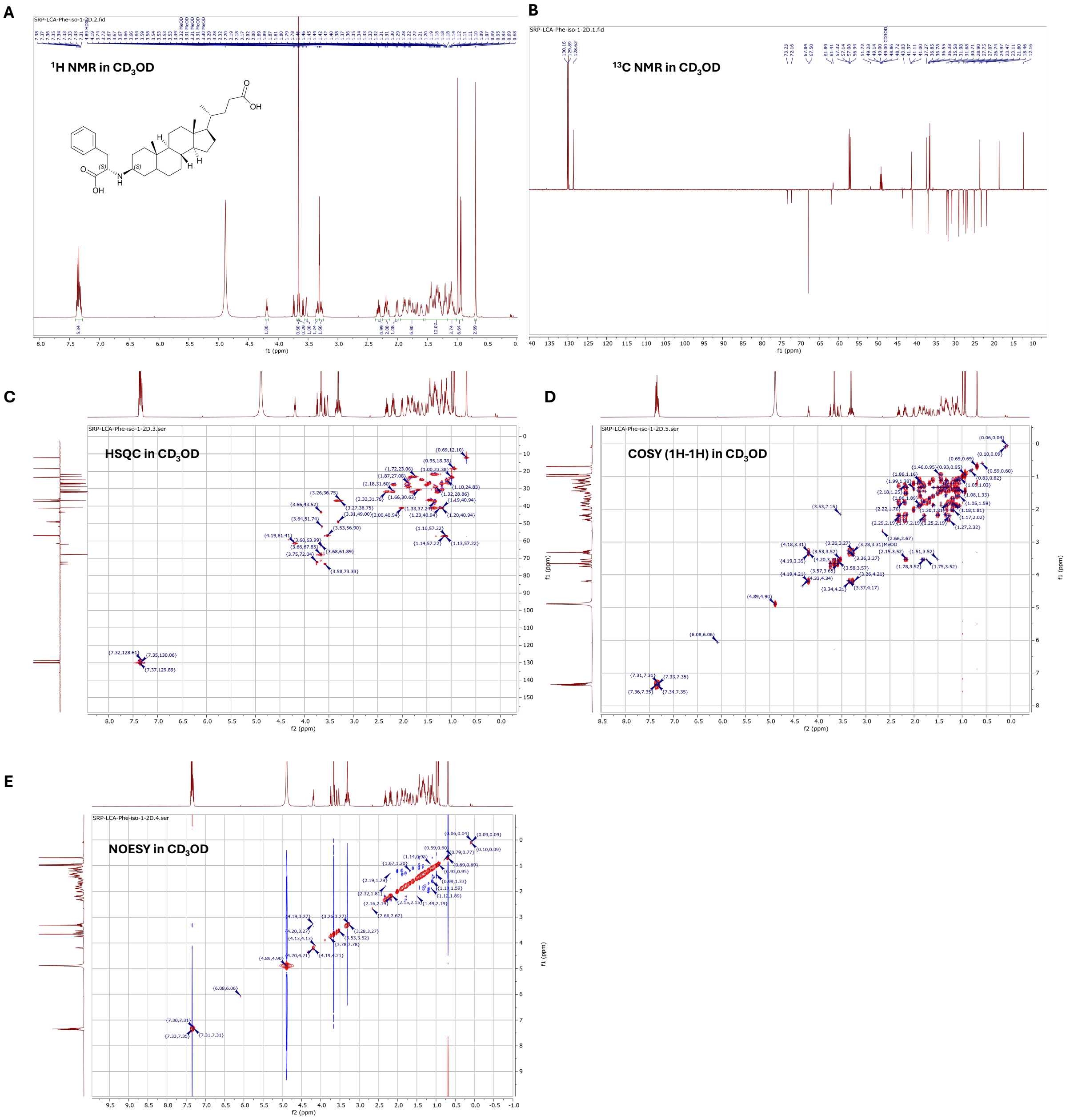


**Fig. S12. 2D NMR characterization of Phenylalanine_3-oxo-lithocholic acid (S) isomer alkamine. The structure of** Phenylalanine_3-oxo-lithocholic acid (S) isomer alkamine was confirmed with, **(A)** $^{1}$H NMR in $CD_3OD$, **(B)** $^{13}$C NMR in $CD_3OD$, **(C)** HSQC in $CD_3OD$, **(D)** COSY (1H-1H) in $CD_3OD$, **(E)** NOESY in $CD_3OD$ using a 600 MHz NMR.

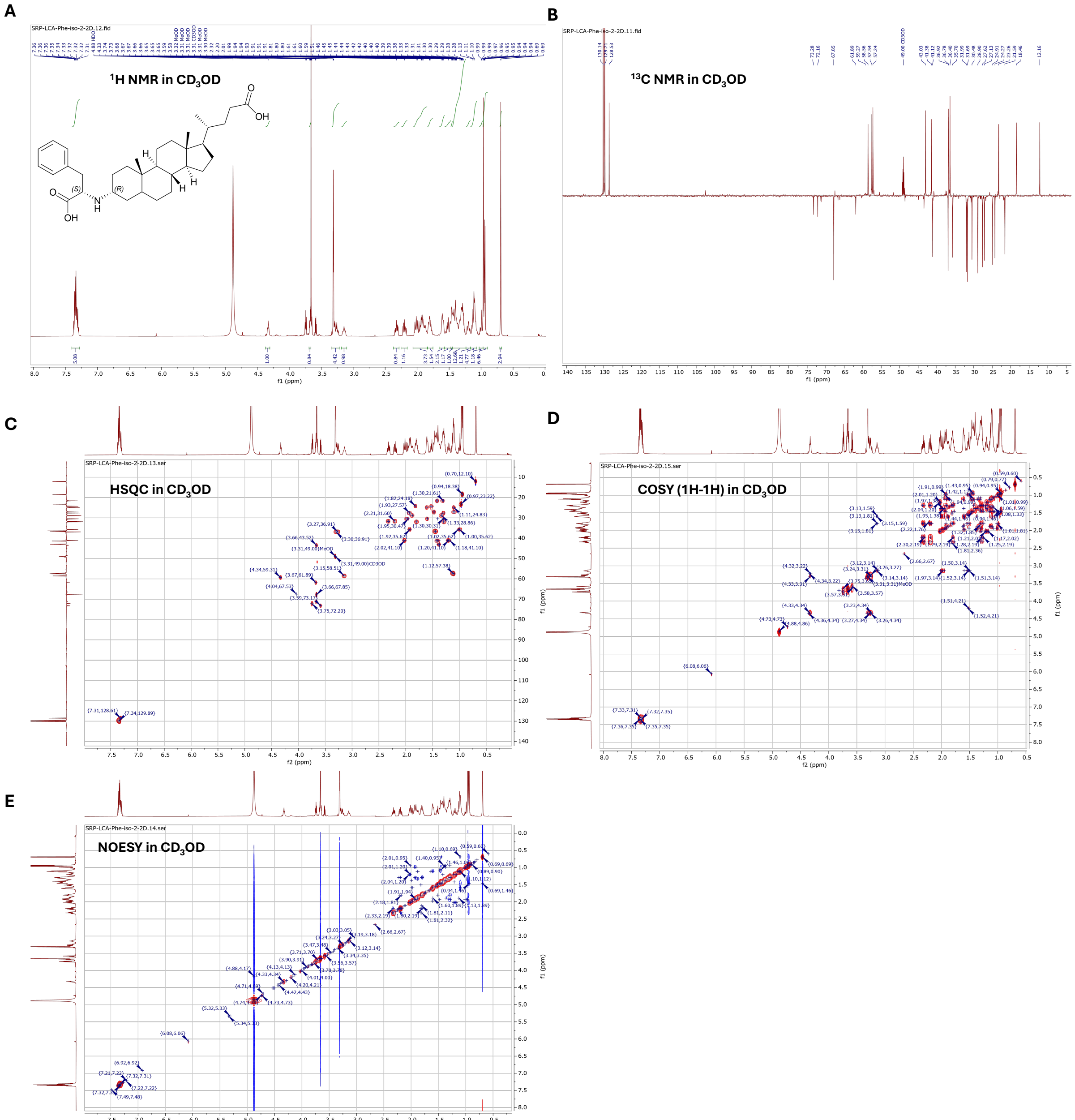


**Fig. S13. 2D NMR characterization of Phenylalanine_3-oxo-lithocholic acid (R) isomer alkamine.** The structure of Phenylalanine_3-oxo-lithocholic acid (R) isomer alkamine was confirmed with, **(A)** $^{1}$H NMR in CD3OD, **(B)** $^{13}$C NMR in CD3OD, **(C)** HSQC in CD3OD, **(D)** COSY (1H-1H) in CD3OD, **(E)** NOESY in CD3OD using a 600 MHz NMR.

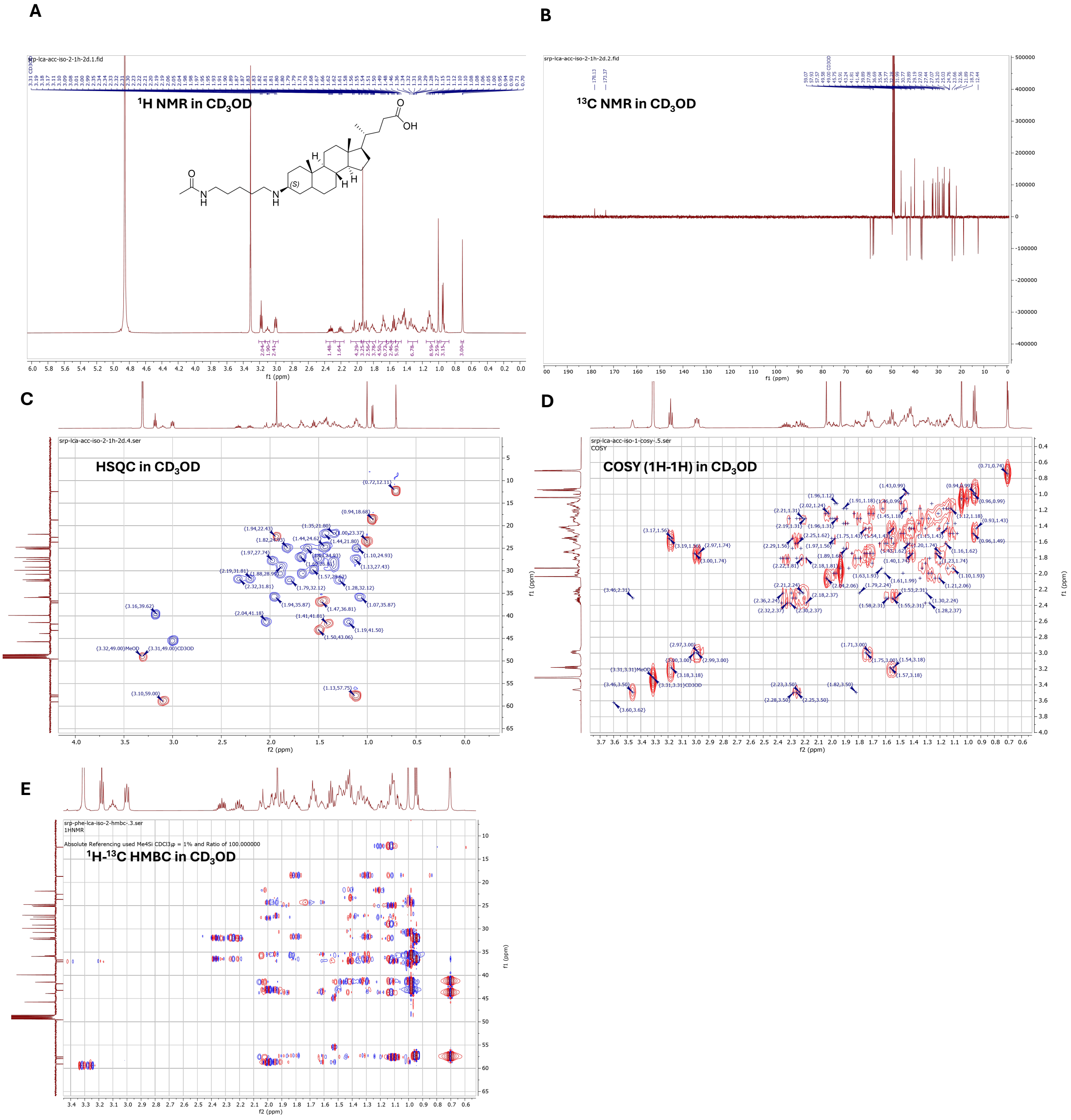


**Fig. S14. 2D NMR characterization of N-acetylcadaverine_3-oxo-lithocholic acid (S) isomer alkamine.** The structure of N-acetylcadaverine_3-oxo-lithocholic acid (S) isomer alkamine was confirmed with, **(A)** $^{1}$H NMR in $CD_3OD$, **(B)** $^{13}$C NMR in $CD_3OD$, **(C)** HSQC in $CD_3OD$, **(D)** COSY (1H-1H) in $CD_3OD$, **(E)** $^{1}$H-$^{13}$C HMBC in $CD_3OD$ a 600 MHz NMR.

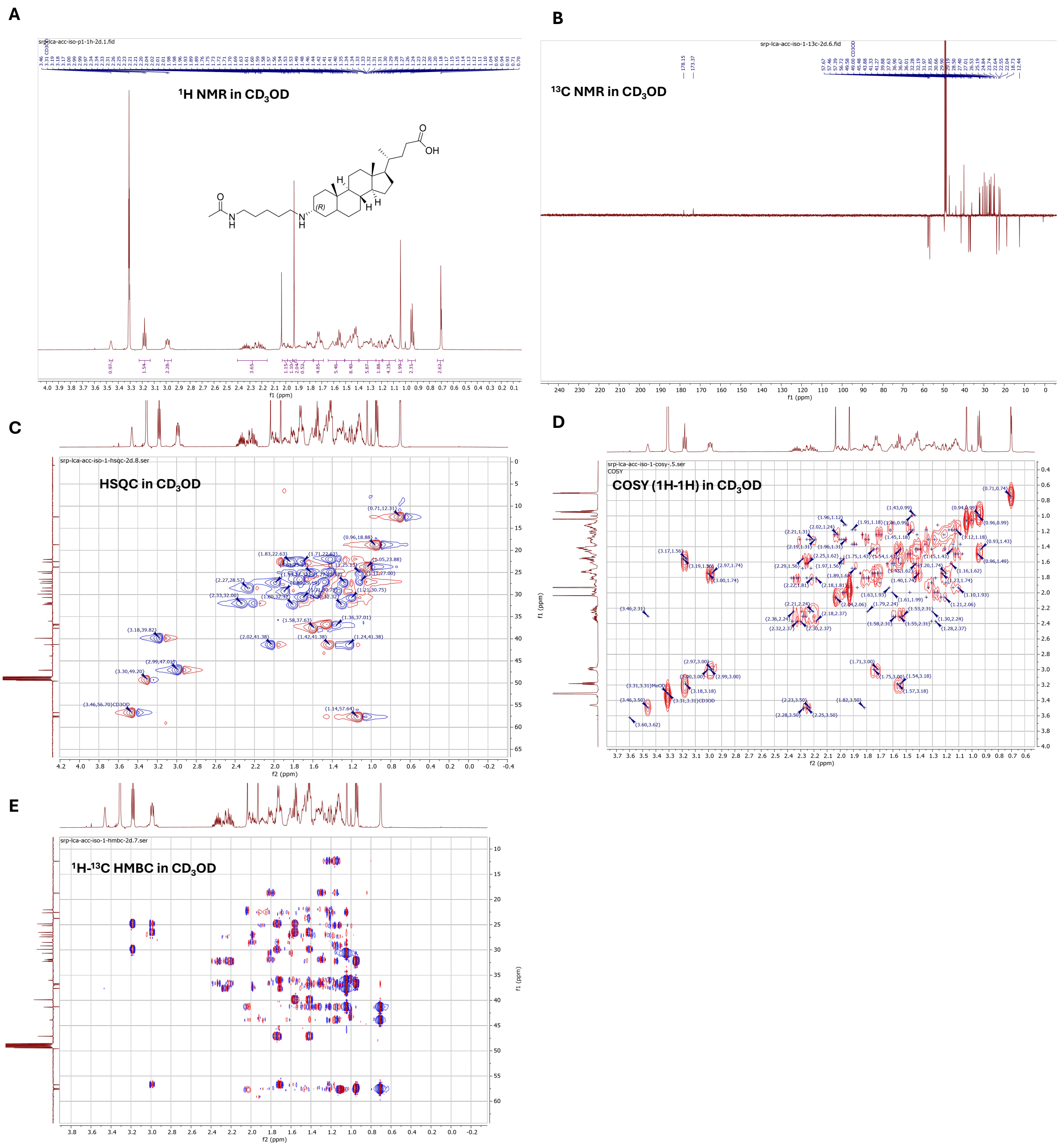


**Fig. S15. 2D NMR characterization of** N-acetylcadaverine_**3-oxo-lithocholic acid (R) isomer alkamine.** The structure of N-acetylcadaverine_3-oxo-lithocholic acid (R) isomer alkamine was confirmed with, **(A)** $^1$H NMR in $CD_3OD$, **(B)** $^{13}$C NMR in $CD_3OD$, **(C)** HSQC in $CD_3OD$, **(D)** COSY (1H-1H) in $CD_3OD$, **(E)** $^1$H-$^{13}$C HMBC in $CD_3OD$ a 600 MHz NMR.

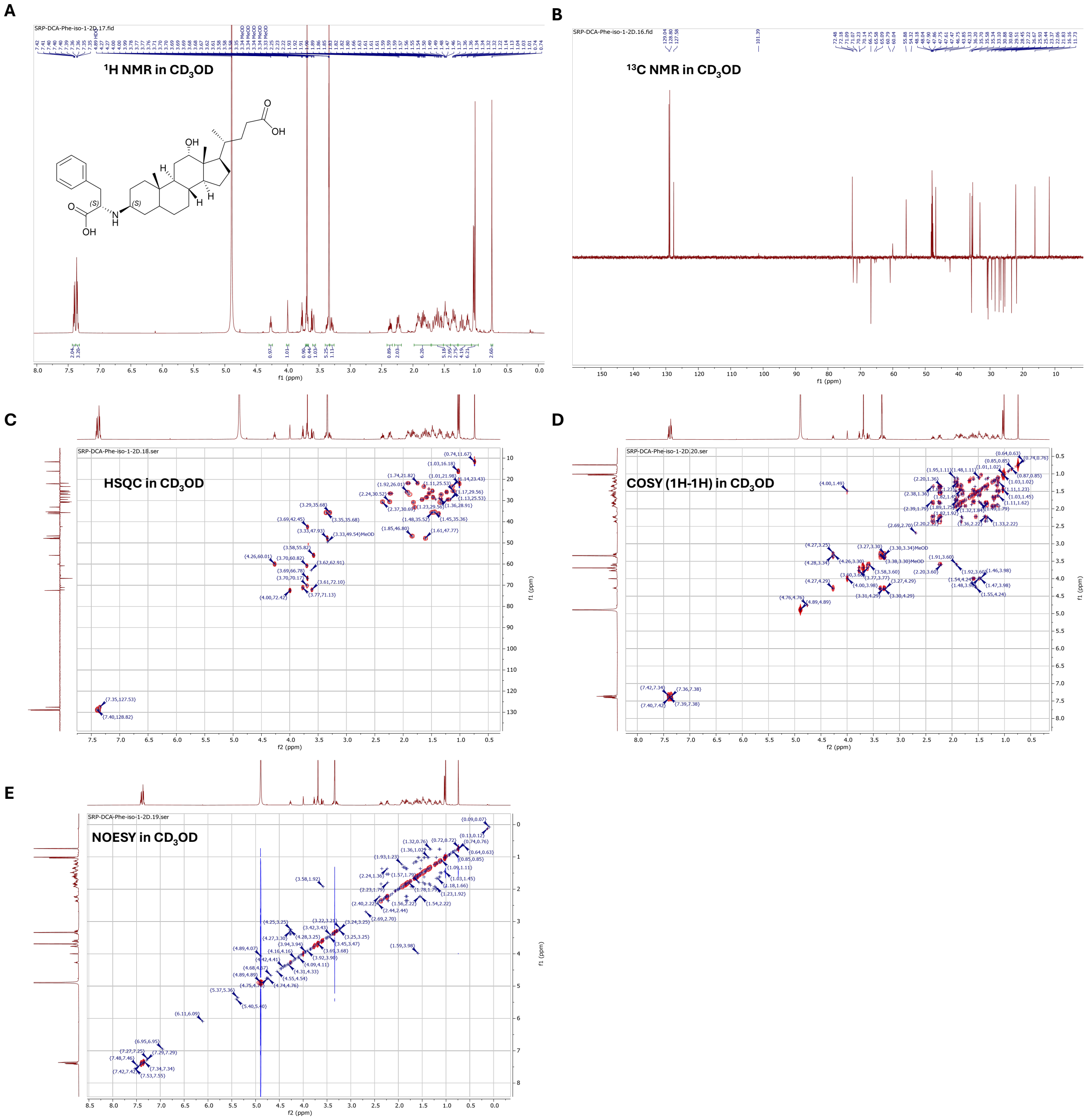


**Fig. S16. 2D NMR characterization of Phenylalanine_3-oxo-deoxycholic acid (S) isomer alkamine.** The structure of Phenylalanine_3-oxo-deoxycholic acid (S) isomer alkamine was confirmed with, **(A)** $^1$H NMR in $CD_3OD$, **(B)** $^{13}$C NMR in $CD_3OD$, **(C)** HSQC in $CD_3OD$, **(D)** COSY (1H-1H) in $CD_3OD$, **(E)** NOESY in $CD_3OD$ using a 600 MHz NMR.

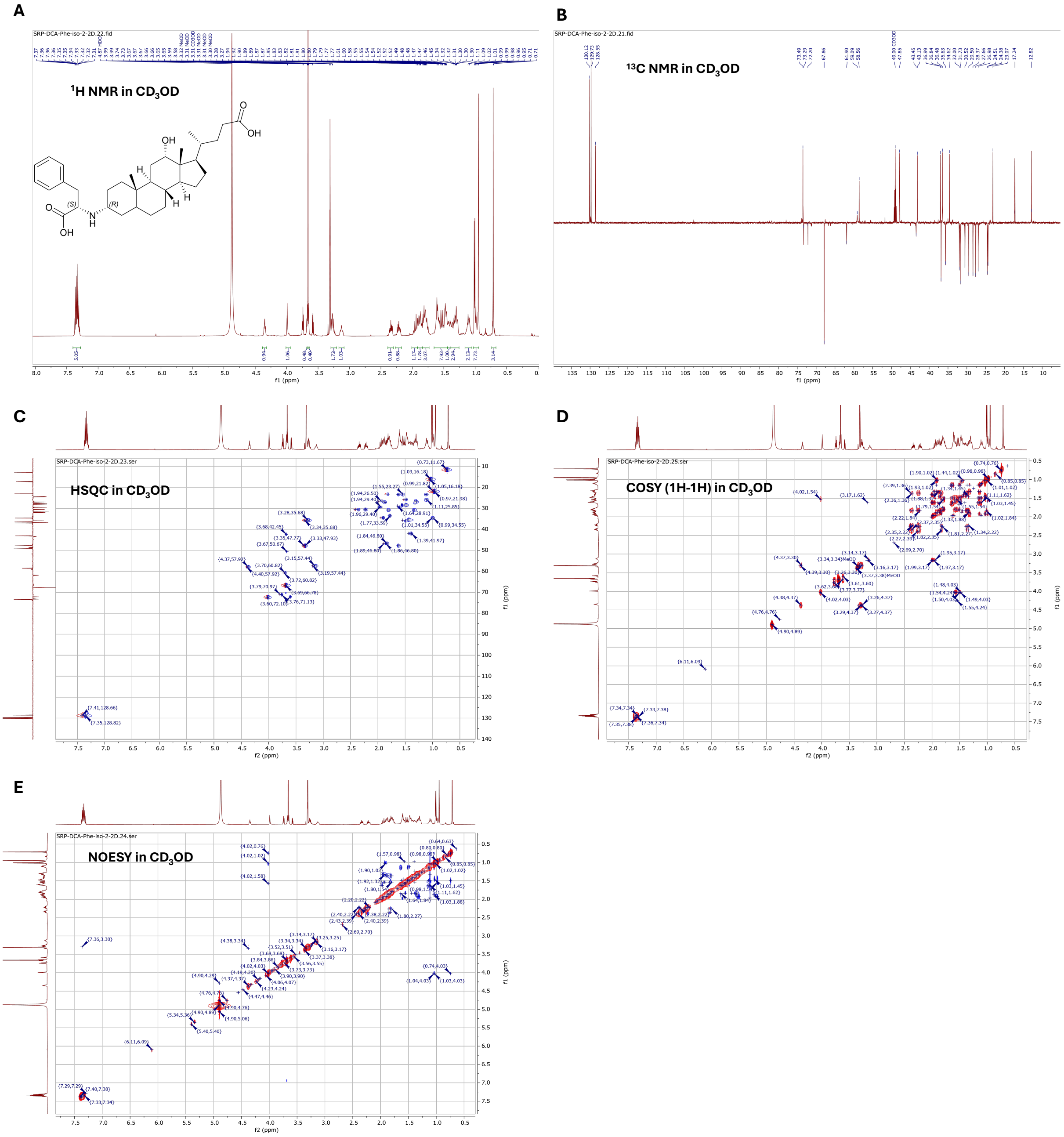


**Fig. S17. 2D NMR characterization of Phenylalanine_3-oxo-deoxycholic acid (R) isomer alkamine. The structure of** Phenylalanine_3-oxo-deoxycholic acid (S) isomer alkamine was confirmed with, **(A)** $^{1}$H NMR in $CD_3OD$, **(B)** $^{13}$C NMR in $CD_3OD$, **(C)** HSQC in $CD_3OD$, **(D)** COSY (1H-1H) in $CD_3OD$, **(E)** NOESY in $CD_3OD$ using a 600 MHz NMR.

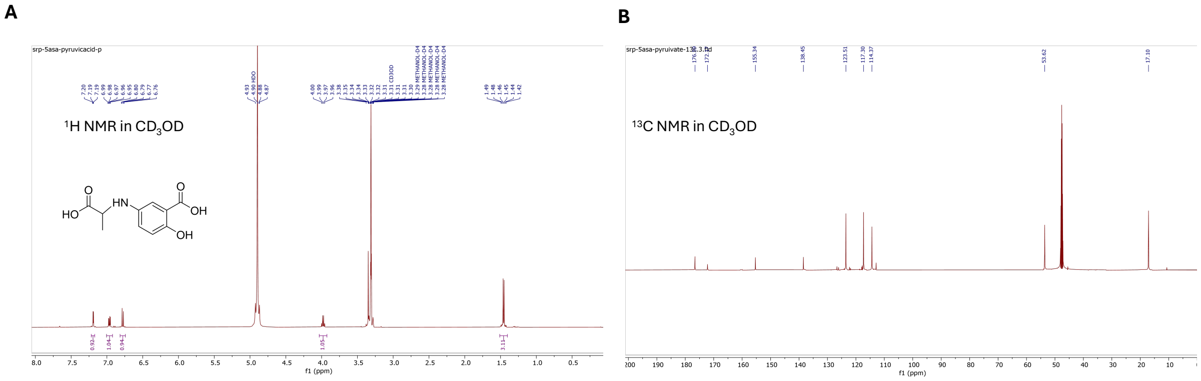


**Fig. S18. 1D NMR characterization of 5-((1-carboxyethyl)amino)-2-hydroxybenzoic acid alkamine. The structure of** 5-((1-carboxyethyl)amino)-2-hydroxybenzoic acid alkamine was confirmed with, **(A)** $^{1}$H NMR in $CD_3OD$, **(B)** $^{13}$C NMR in $CD_3OD$ using 600 MHz NMR.

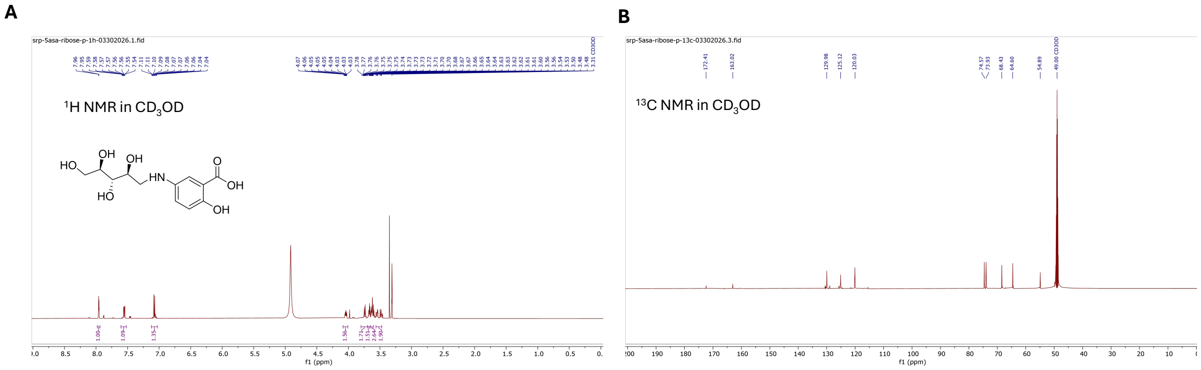


**Fig. S19. 1D NMR characterization of 2-hydroxy-5-(((2S,3S,4R)-2,3,4,5-tetrahydroxypentyl)amino)benzoic acid alkamine. The structure of** 2-hydroxy-5-(((2S,3S,4R)-2,3,4,5-tetrahydroxypentyl)amino)benzoic acid alkamine was confirmed with, **(A)** $^{1}$H NMR in $CD_3OD$, **(B)** $^{13}$C NMR in $CD_3OD$ using 600 MHz NMR.

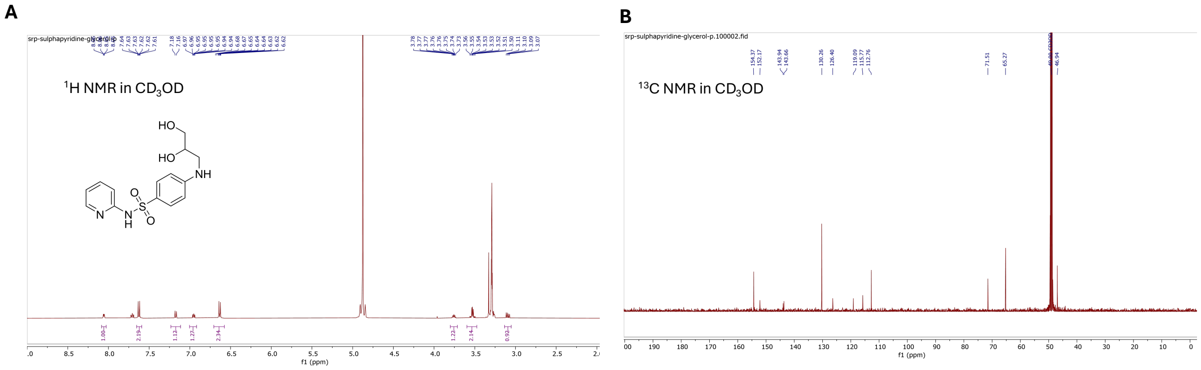


**Fig. S20. 1D NMR characterization of 4-((2,3-dihydroxypropyl)amino)-N-(pyridin-2-yl)benzenesulfonamide alkamine. The structure of** 4-((2,3-dihydroxypropyl)amino)-N-(pyridin-2-yl)benzenesulfonamide alkamine was confirmed with, **(A)** $^{1}$H NMR in $CD_3OD$, **(B)** $^{13}$C NMR in $CD_3OD$ using 600 MHz NMR.

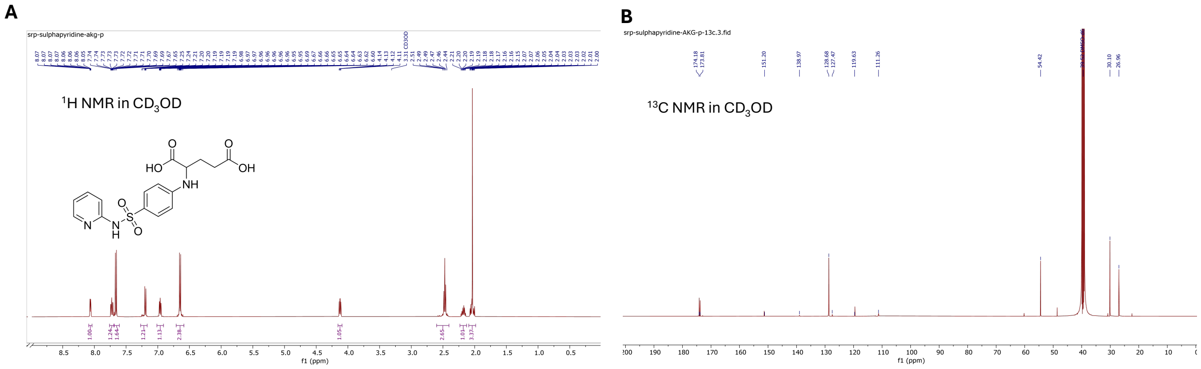


**Fig. S21. 1D NMR characterization of (4-(N-(pyridin-2-yl)sulfamoyl)phenyl)glutamic acid alkamine. The structure of** (4-(N-(pyridin-2-yl)sulfamoyl)phenyl)glutamic acid alkamine was confirmed with, **(A)** $^1$H NMR in $CD_3OD$, **(B)** $^{13}$C NMR in $CD_3OD$ using 600 MHz NMR.
